# Synthetic enhancers reveal design principles of cell state specific regulatory elements in hematopoiesis

**DOI:** 10.1101/2024.08.26.609645

**Authors:** Robert Frömel, Julia Rühle, Aina Bernal Martinez, Chelsea Szu-Tu, Felix Pacheco Pastor, Rosa Martinez Corral, Lars Velten

## Abstract

During cellular differentiation, enhancers transform overlapping gradients of transcription factors (TFs) to highly specific gene expression patterns. However, the vast complexity of regulatory DNA impedes the identification of the underlying cis-regulatory rules. Here, we have characterized 62,126 fully synthetic DNA sequences to bottom-up dissect design principles of cell-state specific enhancers in the context of the differentiation of blood stem cells to seven myeloid lineages. Focusing on binding sites for 38 TFs and their pairwise interactions, we found that identical sites displayed both repressive and activating function, as a consequence of cellular context, site combinatorics, or simply predicted occupancy of a TF on an enhancer. Surprisingly, we found that combinations of activating sites frequently neutralized each other or even gained repressive function. These negative synergies convert quantitative imbalances in transcription factor expression into binary downstream activity patterns, a principle that can be exploited to build differentiation-state specific enhancers from scratch.

## Introduction

During cellular differentiation, the activity of transcription factors (TFs) needs to be transformed to highly specific expression patterns of lineage-specific genes. In the context of blood stem cell differentiation, single-cell transcriptomics^1–3^, as well as imaging-based^4^ studies have demonstrated that key TFs are expressed in smooth, highly overlapping gradients throughout the stem- and progenitor population (Figure 1a, top row); by contrast, the target genes regulated by these factors display cell state-specific expression patterns (Figure 1a, bottom row). Similar overlapping TF gradients accompany cellular differentiation in systems ranging from the exit from pluripotency^5^ over organogenesis^6^ to tissue regeneration by other adult stem cells^7,8^. Indeed, altered ratios between transcription factors, and not their binary combination, can be sufficient to establish specific gene expression programs^9,10^. But how can enhancers transform quantitative imbalances in an identical set of TFs into highly specific activity patterns? How can lineage- and cell cycle genes in stem cells be silenced despite the abundant expression of lineage TFs in these cells? And: How can cancer cells achieve a co-expression of stem- and progenitor programs^11^ while mostly relying on the same core transcriptional regulators?

**Figure 1:**
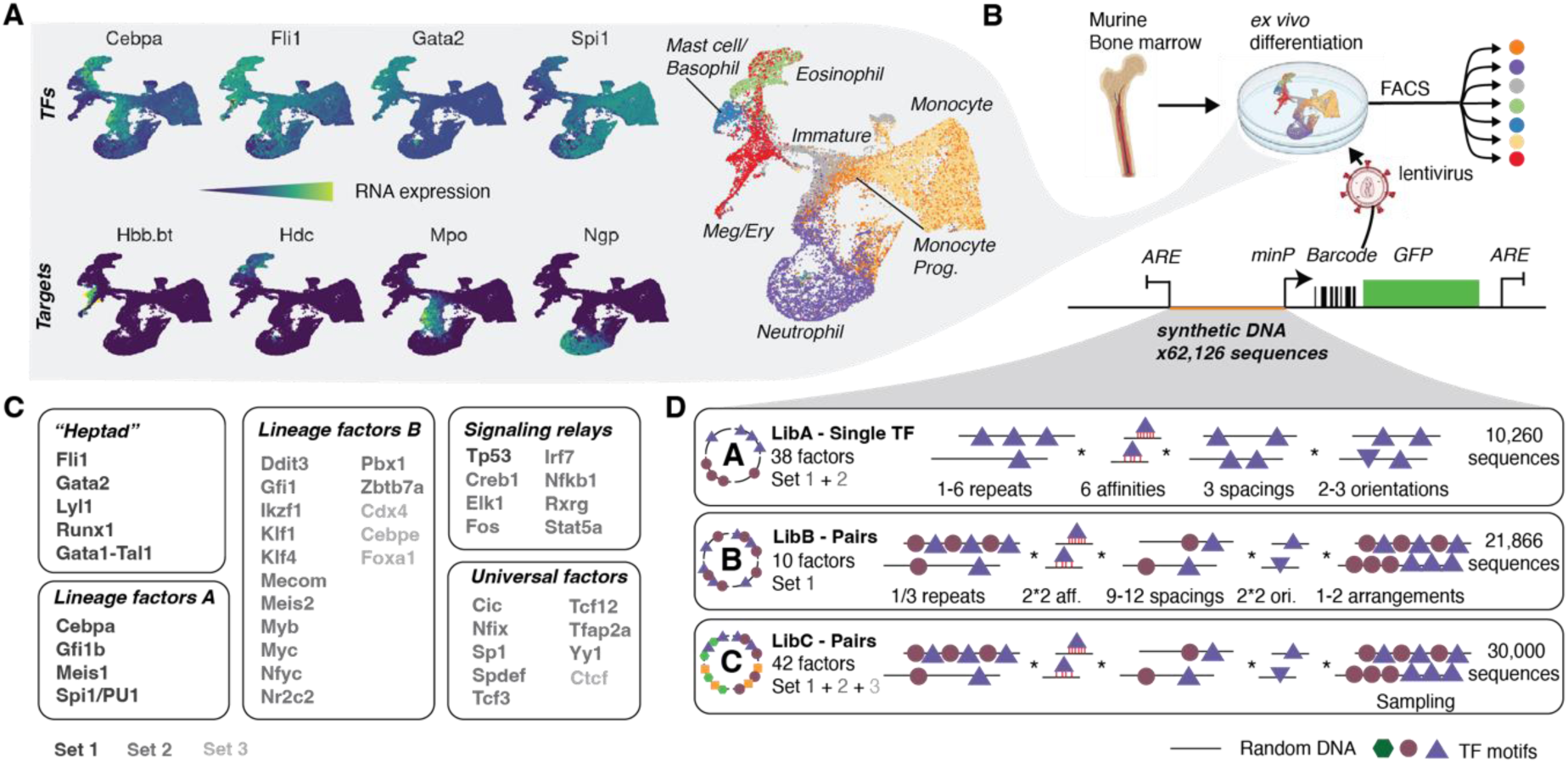
Experimental design. See also Supplementary Note 1 and Figure S1-3. **A.** TFs are expressed in an overlapping manner during early hematopoietic differentiation. uMAPs of single-cell RNA-seq data of hematopoietic progenitor cells in culture highlighting gene expression. Key hematopoietic TFs (top row) are broadly expressed in an overlapping manner whereas the expression patterns of their targets is specific (bottom row). Right panel: Colors indicate cell states purified by FACS, see Figure S2h. **B.** Experimental design. Hematopoietic stem- and progenitor cells were extracted from bone marrow and placed into culture medium supportive of pan-myeloid differentiation^27,29^. Synthetic enhancer constructs were delivered using the lentiMPRA vector^31^, cells were FACS-sorted into seven cell states, and the activity of each synthetic enhancer in each cell state was determined. ARE: Antirepressor Element^30^. minP: minimal promoter. **C.** Selection of TFs. TFs are grouped by functional annotations from literature^14,35–42^. Different sets of TFs (colors) were used for different experiments (see panel D). **D.** Design of candidate DNA libraries. TF motifs (circles/triangles) of single factors (Lib A) or factor pairs (Lib B/C) were embedded in random DNA at different binding site strength, number, and spacing, orientation and arrangement towards each other. See Methods for detail.

To answer these broadly relevant questions, a functional understanding of enhancers and their interactions with TFs is needed. Studies of gene regulatory elements (GREs) from the natural genome are limited in that regard. Genome wide datasets of transcription factor binding and chromatin accessibility^12–14^ are not informative regarding the functional role of each TF in a given context. Experimental manipulations of TF levels^15,16^ or mutagenesis of natural enhancers^17^ demonstrate that single TFs cause non-trivial increases and decreases of the activity of TF-bound enhancers, but the plethora of contexts in which TFs bind make it difficult to understand or predict these effects. Indeed, it has been argued that the natural genome contains only few instances of many transcription factor combinations and binding site arrangements, and is in a statistical sense underpowered to ‘learn the rules by which it is decoded’^18,19^.

An alternative way to study the cis-regulatory code is to measure the enhancer activity of synthetic, systematically designed DNA sequences, where TF bindings sites (TFBS) are placed in an otherwise inert context^17,20–24^. This bottom-up approach keeps strong control over the sequence composition of candidate enhancers, and can provide data of sufficient statistical power to characterize combinatorial action of TFBS, as demonstrated by recent studies that have profiled tens of thousands to millions of synthetic or even random DNA sequences in the context of cell lines or yeast^23,25,26^. There is, however, a paucity of large-scale, functional datasets characterizing systematically designed enhancers during cellular differentiation. In particular, synthetic enhancers have never been used to investigate the cis-regulatory code that transforms overlapping TF gradients to highly cell state specific gene expression patterns. Advanced primary cell models of hematopoietic differentiation^27–29^ provide a unique opportunity to study this broadly relevant question, as they allow to measure the activity of a large number of synthetic DNA constructs in a complex model of stemness and differentiation.

Here, we studied the function and interactions of 38 key TF binding sites during hematopoietic differentiation in minimal synthetic enhancers (Figure 1b-d). Specifically, we placed the binding sites of single TFs, or pairs of TFs, on random DNA in different arrangements, using different motif affinities, and different numbers of repeats (Figure 1d). The ability of 62,126 such constructs to drive transcription was determined by a lentiviral massively parallel reporter assay (lentiMPRA)^30–32^ in seven cell states of pan-myeloid and erythroid murine HSC differentiation^27,29^. Our data allow us to comprehensively characterize the function of binding sites and their pairwise interactions across differentiation. We found that enhancers containing binding sites of individual ‘lineage factors’ (e.g. Spi1, Cebpa, Gata1, Fli1) activate transcription, but specific combinations of these sites are inactive or even act as repressors, creating enhancers that are sensitive to ratios in TF expression and specific to cell states where either factor dominates. Conversely, binding sites of several ‘stem cell factors’ (e.g. Runx1, Meis1) act as repressors, but permit or support activity in specific combinations. Together, these interactions serve to convert overlapping transcription factor gradients into highly cell state specific, binary activity patterns. In the context of a human leukemia cell line commonly used for MPRA studies^33,34^ (K562), by contrast, activators usually dominate and repressive interactions are scarce. In sum, these data highlight negative synergies between TF binding sites as a design principle of cell state specific enhancers in hematopoiesis.

## Results

### Design of synthetic enhancer libraries

We selected 38 key hematopoietic TFs from literature^14,35–42^, as well as from the analysis of scATAC-seq data^43^ (Figure 1c, S1a,b). To systematically characterize the function and interactions of their binding sites, we designed three DNA libraries (Figure 1d, Table S1): Library A investigated the function of binding sites of individual TFs, in the context of random DNA. It contained candidate enhancer sequences with one to six binding sites for one single TF, and systematically explored different options regarding the spacing, arrangement, orientation, number, and affinity of motifs. We refer to each unique combination of these parameters as the ‘design’ of a candidate enhancer. Each factor was covered with 270 candidate enhancers of different designs.

Library B and C investigated the function of pairs of TFs, by placing combinations of binding sites in random DNA. Library B profiled all 45 pairwise combinations of a biologically important set of 10 TFs (Figure 1c: lineage master regulators and ‘heptad’^14^ factors). One or three binding sites for either factor were placed on random DNA while options for the spacings, orientations and arrangements of motifs were systematically explored. Library C profiled a total of 861 factor combinations, covering each pair with 22-34 candidate enhancers sampled from the complex space of possible designs. Besides the 38 core TFs from Library A, it also included 4 further TFs with relevant effects in recent genome-wide MPRA data (ref. ^33^ and own analyses). For the design of the enhancers, motif instances of a defined affinity were sampled from a consensus PWM (Figure S1c-f, methods), and placed on random background DNA filtered to avoid strong binding sites of the 38 factors. Each candidate enhancer we assayed had a different random DNA background, and all motif instances were independently sampled. Thereby, we made sure that our libraries were not biased towards a specific DNA context or motif instance.

Libraries were cloned in front of a minimal promoter in a lentiviral MPRA reporter vector^30,31^. In lentiviral MPRA, enhancer activity is determined after integration into the genome, i.e. DNA is covered in histones. Primary mouse hematopoietic stem- and progenitor cells (HSPCs, defined as Lin^-^Kit^+^) were infected with the MPRA libraries and allowed to differentiate for four days, using conditions^27,29^ that support the parallel differentiation into a heterogeneous population of hematopoietic progenitors (HPCs), including precursors of red blood cells, megakaryocytes, neutrophils, eosinophils, basophils, and monocytes. We characterized these cultures using single-cell RNA-seq and CITE-seq, confirming that they recapitulate gene expression patterns observed *in vivo* (Figure 1a, Supplementary Note 1, Figure S2a-d). We then designed and validated a FACS gating scheme to sort the cultures into seven well-defined progenitor cell states (Figure 1a, Figure S2e-g) and measured the activity of the MPRA constructs in these seven cell states. Additionally, all libraries were also screened in K562 cells derived from a chronic myeloid leukemia patient. In total, 398,324 cell-state specific measurements of GRE activity in HPCs passed our quality filters; each of these measurements was based on averaging 50-100 lentiviral integration sites per construct, i.e. integration site effects were efficiently averaged out (Figure S3c). In K562s, we obtained 104,717 measurements based on averaging 100-1000 lentiviral integration sites per construct (Figure S3d). Correlations between replicate infections were 0.95-0.99 in the cell line, and 0.74-0.87 in primary cells (Figure S3e,h,i). A detailed characterization of our primary cell MPRA setup is provided in Supplementary Note 1 and Figure S2, S3.

In sum, our synthetic DNA libraries explored the cell state specific activities of synthetic enhancers composed of binding sites for single TFs and pairs of TFs in the context of random DNA. The relationship between sequence design (number and affinity of binding sites, spacing between sites, site orientation, site arrangement) and resulting activity can be explored interactively for all constructs in our web-based resource, http://veltenlab.crg.eu/MPRA.

### Combinatorial interactions between TFs establish binary activity patterns

We first analyzed the data from Library A and B qualitatively, to characterize the predominant effect of factors and factor pairs as activators or repressors of transcription in each cell state. To that end, we made use of several hundred random DNA sequences included in all libraries that served to establish a baseline range of activities (defined as zero). We calculated the 5^th^ and 95^th^ percentiles of the activity of these sequences to define repression and activation thresholds, respectively (Figure 2a), and classified each candidate enhancer as “active” if it was beyond the activation threshold, and “repressed” if it was below the repression threshold (Figure 2a). We then determined for each TF in each cell state the fraction of sequences containing its motif that are either active or repressed, and determined if that fraction exceeded the expectation from random DNA by a resampling-based statistical test.

**Figure 2:**
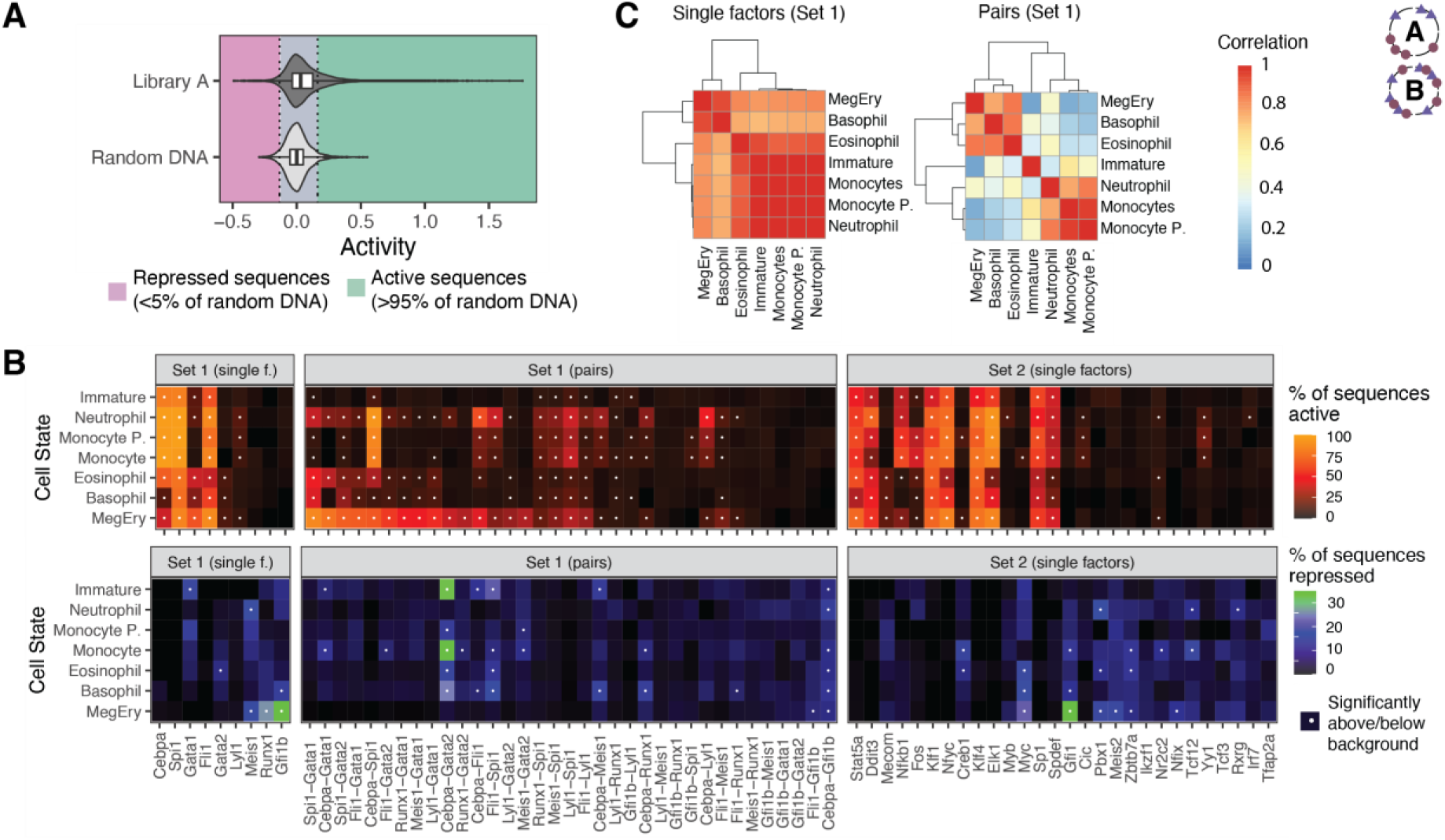
TF pairs, but not single factors, activate gene expression in a lineage-specific manner. Analyses are based on Library A and B, see Figure 1d for design. **A.** Identification of active and repressed sequences. Violin plot illustrating the activity of random DNA and sequences from Library A. Regions corresponding to activity smaller than the 5^th^ percentile of random DNA, or bigger than the 95^th^ percentile of random DNA, are color coded. **B.** Activity of factors and factor pairs across seven cell states. Heatmaps depicting for all factors or factor pairs from Set 1, the fraction of sequences containing the respective motif(s) that are active (upper row) or repressed (bottom row) in each of the seven cell states. White dots indicate that activation/repression strength is larger than expected for random DNA, based on a resampling based statistical test. Trp53 and its combinations are excluded because they are always active, see Figure 5a. **C.** Activity patterns of enhancers containing factor pairs, but not single factors, separate lineages. Heatmap depicting correlation in factor activity, as determined in panel B, across lineages.

We thereby found that most sequences containing binding sites for single factors were either generally inactive across cell states, or active in most cell states (Figure 2b, top row). Evidently, some factors were mostly active in erythromyeloid lineages or in monocytes/neutrophils; however, no single factor was able to strongly drive expression exclusively in a single lineage or sets of lineages. Repression was more specific (Figure 2b, bottom row). By contrast, factor pairs induced binary patterns of activity, inactivity and repression (Figure 2b). In particular, many combinations of factors were able to activate transcription in some cell states, and repress it in others.

The ability of single factors to activate transcription was highly correlated between all lineages, and only roughly separated lineages into an erythromyeloid and a myeloid branch (Figure 2c). By contrast, factor pairs were only correlated between closely related lineages and separated lineages hierarchically according to the known transcriptional and developmental relationships between them^3^ (Figure 2c). Taken together, binding site pairs can establish a binary activity pattern of enhancers across a differentiation landscape characterized by highly overlapping TF expression patterns, whereas enhancers composed of single motifs can in most cases only establish gradual differences.

### Enhancers as continuous relays of TF concentration

To dissect these phenomena into design rules of lineage-specific enhancers, we first focused on Library A, designed to investigate the regulation of gene expression by single factors. Across the seven cell states, 7/38 factors were inactive in all cell states, 22/38 were exclusively activators of transcription, and 7/38 were exclusively repressors (Gfi1b, Gfi1, Runx1, Meis1, Meis2, Pbx1 and Tcf12) (Figure 3a, Table S2). Some of these factors have previously been described as repressive in the context of hematopoiesis^36,44^. In addition to activators and repressors, we found two factors (Mecom and Myc) that were repressive in some cell states, and activating in others (Figure 3a, Figure S4a). We refer to this ability of a factor to exert repression and activation in a cell-type specific manner as *cell context dependent duality*.

**Figure 3:**
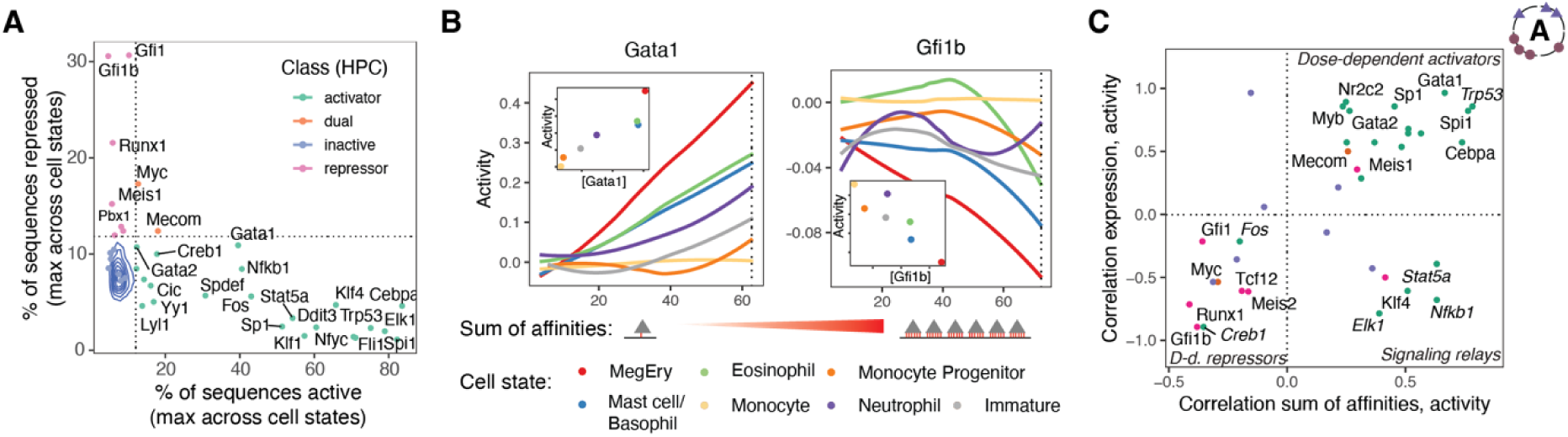
Characterization of single TFs as activators, repressors or dual factors. Analyses are based on Library A, see Figure 1d for design. See Table S2 for figure source data. **A.** Identification of activators, repressors, inactive and dual factors. Scatter plot determining, for each factor, the fraction of sequences containing its binding site that are more active, compared to the 95^th^ percentile of random DNA (x-axis) and the fraction of sequences that are less active, compared to the 5^th^ percentile of random DNA (y-axis). The maximum active/repressed percentage across the seven cell states is shown. Blue lines indicate probability density from a resampling-based analysis of random DNA. Dashed lines indicate upper bound of a 95% confidence interval. Colors group factors into activators, repressors or dual factors based on crossing that bound. **B.** Relationship between enhancer design, TF expression, and enhancer activity. Main panels: Line plots displaying smoothened means of activity, conditioned by the sum of motif affinities contained in the sequence and stratified by cell state, for Gfi1b and Gata1. See Methods, section *Affinity computation and data visualization.* Insets: Scatter plots comparing factor expression levels from RNA-seq (x-axis) the activity at the dashed line (y-axis). **C.** Scatter plot depicting, for each factor, the correlation between sum of motif affinities and activity in the cell state with the highest activity for a given factor (x-axis), and the correlation between factor expression and the activity, at the highest sum of motif affinities (y-axis). Factors are color-coded by class, see panel A for a color code.

Across the seven cell states, activity levels were positively correlated with mRNA expression of the corresponding transcription factor in the case of 16/22 activators, and negatively correlated in the case of 5/7 repressors (Figure 3b, insets, and Figure 3c, y axis). Known signaling relays subject to post-translational regulation (Stat5a, Nfkb1, Elk1) did not display positive correlations between mRNA expression of the factor and activity of sequences containing its motifs. Further exceptions were Klf4 and Gfi1, where activity was instead correlated with the RNA expression of the closely related factors Klf1 and Gfi1b (Figure S4b). These analyses confirm that transcription factor binding sites placed in synthetic DNA respond to changes in the concentration of the corresponding TF.

To further characterize the effect of local TF dose on enhancer activity, we exploited that by design, the synthetic GREs from Library A contained between one weak and six strong binding sites. We found that for 20/22 activators, more (and/or stronger) binding sites led to a higher activity, whereas for 5/7 repressors, more (and/or stronger) binding sites induced stronger repression below baseline (Figure 3b, main panels, and Figure 3c, x axis, Table S2). The relationship between binding site number/affinity and activity was always continuous, and we did not observed all-or-nothing responses.

Together, these analyses classify factors as activator or repressors in HPCs. They demonstrate that enhancers composed of single factors mostly relay local TF concentration to a graded and monotonous activity response, but they also describe a first type of activator-repressor duality, *cell context dependent duality*.

### Occupancy-dependent duality: Enhancers as bandpass filters

For some single factors, however, we observed non-monotonous relationships between binding site number/strength and activity. For these factors, enhancers containing an intermediate number of motifs or motifs of intermediate affinity were maximally activating (Figure 4a). A statistical analysis identified monotonously increasing relationships for 16/22 activators, monotonously decreasing relationships for 4/7 repressors, and non-monotonous relationships for seven factors (Figure 4b).

**Figure 4:**
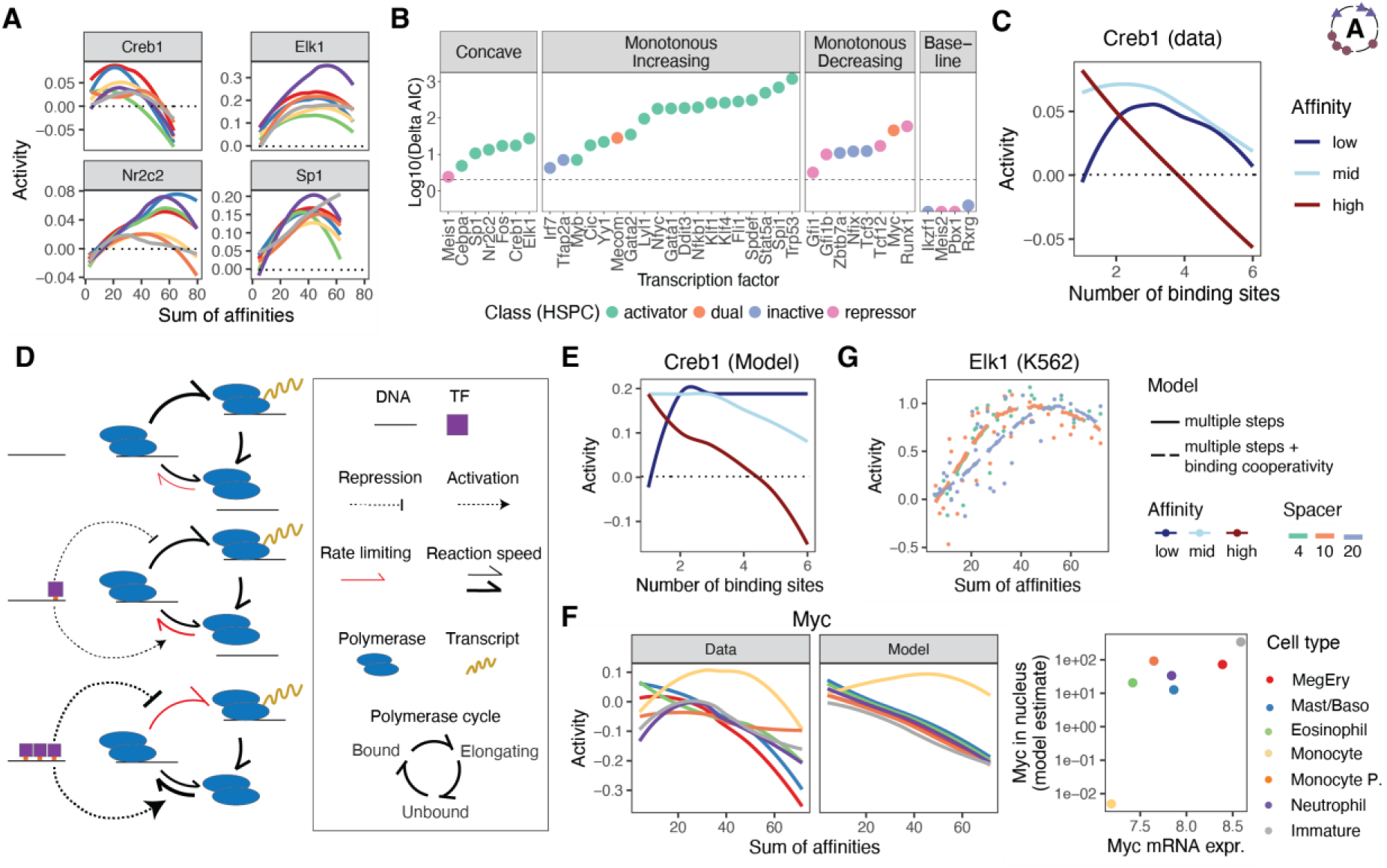
Non-monotonicity and occupancy-dependent duality enable enhancers to act as band pass filters. Analyses are based on Library A, see Figure 1d for design. **A.** Examples of non-monotonous behavior. Line plots displaying smoothened means of activity, conditioned by the sum of motif affinities contained in the sequence, for Creb1, Elk1, Nr2c2 and Sp1. Colors correspond to cell states, see panel F for a color scheme. **B.** Statistical evaluation of non-monotonicity. The relationship between sum of motif affinities and activity (see panel A) was fitted with splines of different constraints (flat line as baseline model, monotonically increasing, monotonically decreasing, or concave) for each factor, using cell state as a covariate. Factors were grouped by the best model (facets). Y-axis depicts the difference in Akaike’s information criterion between the best model and the second-best model. See methods, section *Machine learning based analyses* for detail. **C.** Dissection of non-monotonous behavior for Creb1. Line plots displaying smoothened means of activity, conditioned by the number of motifs and stratified by motif affinity, for Creb1. **D.** Illustration of a model where TFs bind to DNA and activate one step in a polymerase cycle while repressing another step. See methods, section *Biophysical models*. **E.** Line plots displaying model fits for Creb1. **F.** Left panel: Illustration of model and data for Myc. Right panel: Scatter plot depicting the relationship between Myc mRNA expression, and the model estimates of Myc nuclear concentration, across cell states. The only difference across cell states in the model is the concentration of Myc. **G.** Data (points) and model fits (lines) for Elk1. The model considers the effect of space on modulating binding in a phenomenological manner, allowing for a space-dependent concentration parameter, but not space-dependent cooperativity parameters (Methods).

For three of these non-monotonous factors (Creb1, Elk1 and Sp1), we identified a particularly interesting relationship between design and activity: One or two strong binding sites resulted in activation, but more strong sites reduced activity or even resulted in (weak) repression. By contrast, several weak binding sites were needed to induce activation, which turned into inactivity when a maximum number of sites were placed (Figure 4c, S4c). These observations can quantitatively be explained by a biophysical model^45^ where a TF activates a rate limiting step in a transcriptional cycle, and represses other steps, such that a second step becomes rate limiting once a certain TF occupancy is exceeded (Figure 4d,e, see Methods: *Biophysical models*). This model can also explain the cell-context-dependent duality observed for Myc, which required low expression levels to achieve an occupancy sufficiently low to drive activation (Figure 4f). Hence, some single transcription factors can achieve binary activity patterns on their own, by acting as an activator at low occupancy, and as a repressor at high occupancy. Enhancers composed of binding sites for these factors turn into bandpass filters of TF activity.

In the case of Elk1, the ‘turning point’ between activation and repression was further modulated by the spacing between sites, in line with a model where TFs bind more efficiently at a specific spacing, leading to a higher local factor concentration at the same number of binding sites (Figure 4g). This example demonstrates that relatively simple mechanisms can give rise to non-trivial behavior of synthetic DNA elements containing binding sites for single factors.

In sum, these analyses characterize a second type of activator-repressor duality, *occupancy dependent duality*, that can turn an enhancer into a bandpass filter of TF concentration.

### Combinatorial duality: Pairs of activators turn into cell state specific repressors

Combinatorial interactions between motifs are key to transform TF gradients into specific activity patterns. We therefore focused on Library B, which included pairwise combinations of sites for 10 TFs with central importance in hematopoiesis (Figure 1c). Of these, Runx1, Meis1, Gata1/Tal1, Gata2 and Lyl1 are part of the ‘heptad’ of transcription factors with frequent co-binding to hematopoietic stem cell enhancers^14^, and Gata1/Tal1, Fli1 and Cebpa are key master regulators of lineage differentiation.

We quantified each pair’s ability to induce repression or activation beyond the thresholds defined by random DNA across cell states, and surprisingly found that several pairs of activators (Cebpa-Gata1, Cebpa-Gata2, Cebpa-Fli1 and Fli1-Spi1) were repressors in some cell state and activators in others (Figure 5a,b). Importantly, enhancers that contained these motifs individually did not elicit substantial repression in any cell state (Figure 5a,b, see also Figure 3a). These cases therefore constitute a third case of activator-repressor duality, *combinatorial duality*.

**Figure 5:**
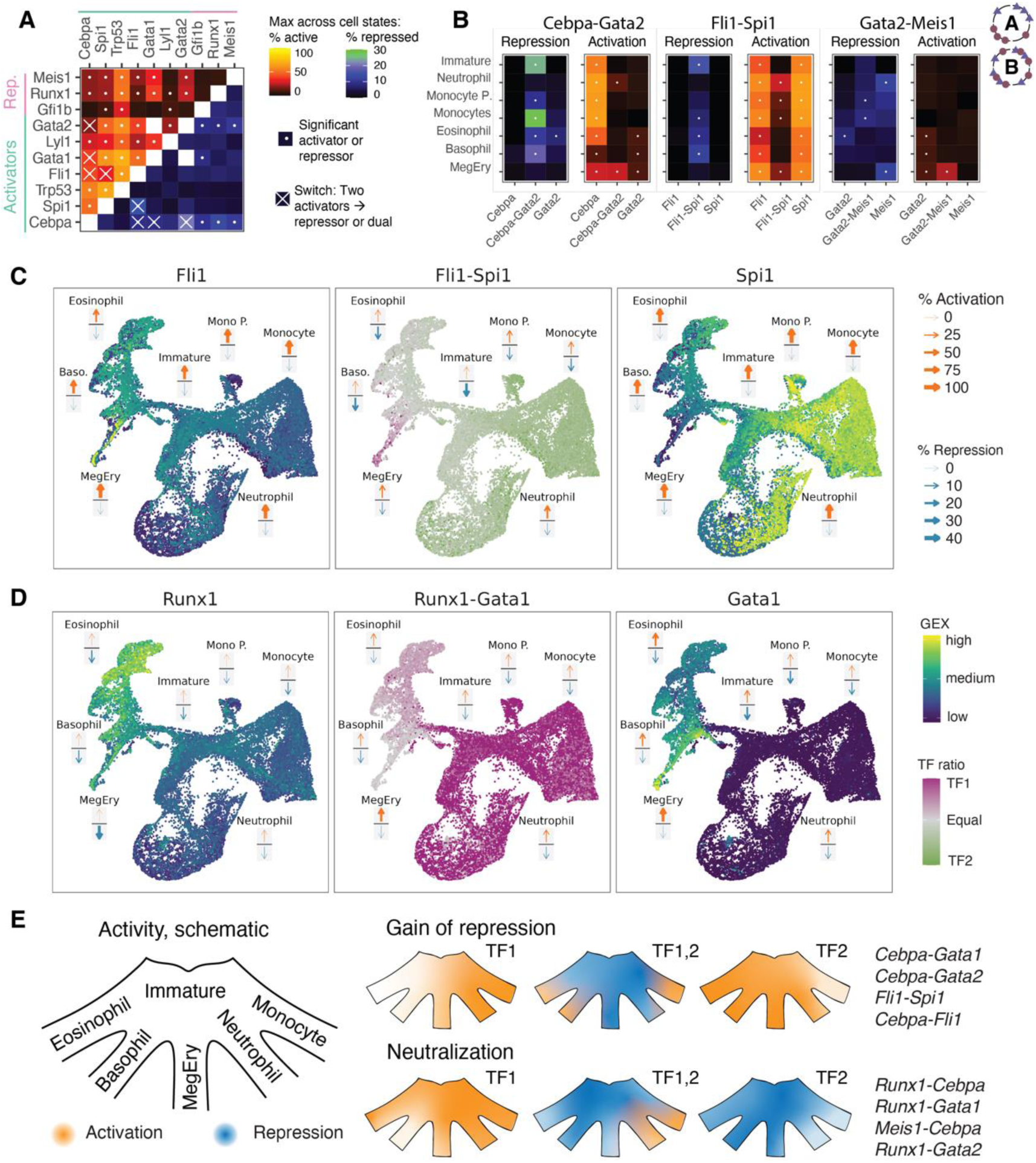
Negative synergies enable enhancers to sense transcription factor ratios. Analyses are based on Library A and B, see Figure 1d for design. **A.** Specific pairs of activators turn into repressors. Heatmap indicating, for each factor pair, the maximum fraction of sequences active or repressed across lineages. Dots indicate that a factor combination was classified as activator (upper triangular matrix) or repressor (lower triangular matrix). Crosses indicate cases where two activators turned into a repressor or dual factor. **B.** Examples of activators turning into repressors. For three factor pairs, heatmaps depicting the fraction of active or repressed sequences in the seven cell states. See panel A for color sale. **C.** Relationship between gene expression and enhancer activity. uMAPs depicting gene expression of Fli1 (left panel), Spi1 (right panel) and the ratio between the two factors’ expression (central panel). Arrows superimposed on the uMAP indicate activation and repression of sequences containing binding sites for Fli1 (left panel), Spi1 (right panel) or both factors (central panel) in the seven cell states. See also Figure 1a for annotation of the uMAP. **D.** Like panel C, but for Runx1 and Gata1. **E.** Scheme illustrating the concept that gradients in transcription factor activity get transformed to highly lineage specific enhancers activity by pairs of activators that turn into repressors, and only allow activity in cell states where either factor dominates.

While Gfi1b was a repressor that dominated over most activators, Runx1 and Meis1 individually were repressors, but displayed specific neutralizing interactions with other TFs. Meis1, together with Gata2, became a very specific activator in megakaryocyte-erythroid precursors, and a repressor in other cell states (Figure 5b). Similar behavior was also observed for the combination of Runx1 and Gata2 (Figure S4d). Cebpa-Runx1 and Cebpa-Meis1 were active very specifically in neutrophil progenitors, and inactive or repressed elsewhere (Figure S4d). Runx1 has previously been described as a specific activator of transcription in context with GATA factors^46^ and a repressor in other contexts^44^.

Together, these analyses demonstrate that the function of hematopoietic TFs as activators or repressors is highly dependent on co-binding TFs, and that interactions with Runx1/Meis1/Fli1 increase the lineage specificity of Gata1/Gata2/Cebpa/Spi1. In particular, we observe several cases of *combinatorial duality*, where a combination of activators turns into a repressor, as well as neutralizing interactions between TFs.

### Combinatorial enhancers sense TF ratios

To relate this functional behavior to TF expression levels, we developed a visualization that superimposes cell state specific patterns of activation and repression (Figure 5c, arrows) on a gene expression landscape measured by single cell RNA-seq (Figure 5c, uMAP). In the case of the Fli1-Spi1, this demonstrated that both factors were expressed over the entire landscape, and activated enhancers containing either individual binding site in all cell states (Figure 5c, left and right column). By contrast, enhancers containing both binding sites were only active in Meg/Ery, Neutrophil progenitors and monocytes, where the balance of the two factors’ expression was imbalanced in favor of either factor (Figure 5c, central column). A similar behavior was observed for Cebpa-Fli1 and Cebpa-Gata2 (Figure S5a,b).

Neutralizing interactions of Runx1/Meis1 with Cebpa/Gata1 (Figure 5d, Figure S5c,d) equally acted as sensors of TF ratios: Gata1 and Runx1 were both expressed in the entire erythromyeloid branch, and acted as an activator and repressor throughout this branch, respectively. In Meg/Ery progenitors, Gata1 dominated, and combinatorial enhancers became activating. In eosinophil and basophil progenitors, both factors were present, and combinatorial enhancers were inactive (Figure 5d).

Together, these examples demonstrate that negative synergies between TF motifs enable enhancers to sense TF ratios and thereby activate gene expression in a highly cell state specific manner. Our data draw a picture where single TFs are broadly expressed, and broadly activating, across the hematopoietic differentiation landscape. By contrast, pairs of TFs act as repressors or are inactive, except in specific cell states characterized by an imbalance in TF expression (Figure 5e). This principle resulted in synthetic enhancers that are highly cell state specific.

### Interactions giving rise to combinatorial duality depend on enhancer grammar

To gain first mechanistic insights into these interactions, we investigated their dependency on the spacing and orientation of binding sites in our libraries. Specifically, we used a random forest machine learning model to identify TFs and TF pairs for which motif ‘grammar’, i.e. the spacing between motifs and their orientation, had a substantial effect on activity. We found that TF interactions that turn pairs of activators into repressors were dependent on spacing between sites and the orientation of the sites towards each other (Figure 6a, S6a). In the example of Cebpa-Gata1, repression was achieved by sites that were close to each other, whereas more distantly located sites were permissive of transcription (Figure 6b). A similar observation was made for Fli1-Spi1 and Fli1-Cebpa, where strong Fli1 binding sites in close vicinity to Spi1/Cebpa sites induced repression or inactivity (Figure 6c,d). Several other interactions (Fli1-Runx1, Gata2-Gfi1b, Trp53-Gata1) were also highly dependent on the spacing between sites and their orientation. Together, these analyses suggest that specific protein-protein interactions between these TFs or their co-factors underlie the switch of activators to repressors. Similar analyses in the context of Library A identified factors that are known to preferentially bind to DNA as multimers (Stat5a, Nfkb1, Trp53, others; Figure S6b-d).

**Figure 6.**
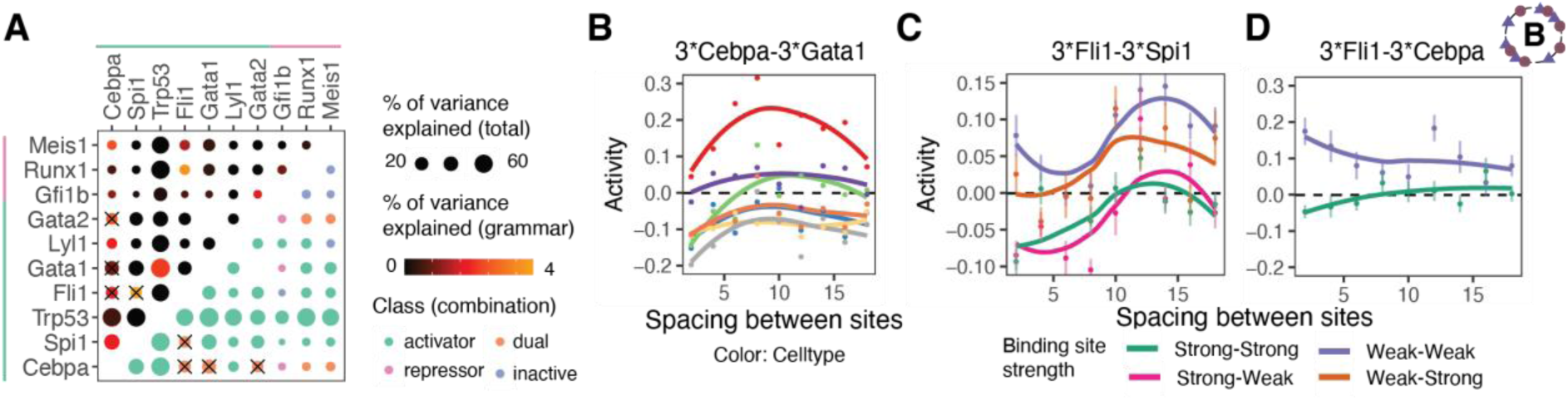
Negative synergies are modulated by enhancer grammar. Analyses are based on Library B, see Figure 1d for design. **A.** Identification of grammar-dependent factor interactions. For each pair of TFs, random forest models were trained on all features of design (i.e. binding site number, binding site affinity, binding site arrangement, spacing between motifs and orientation of motifs) and the performance in ten-fold cross validation was compared to models trained on binding site number and affinity only. In the upper triangle, point size indicates performance of the complete model, color indicates difference between the models. The lower triangle of the matrix classifies factors into activators, repressors or dual factors. **B.** Examples of grammar-dependent factor interactions. Line plot displaying smoothened means of activity, conditioned by the spacing between motifs, stratified by cell state, for Gata1-Cebpa. Only sequences with three binding sites for both factors at alternating arrangement are included in the analysis. **C.** Line plot displaying smoothened means of activity, conditioned by the spacing between motifs and stratified by motif affinities, for Fli1-Spi1 in cell states where this combination elicits repression. Only sequences with three binding sites for both factors are included in the analysis. **D.** Like panel C, except that data for Fli1-Cebpa is shown.

In sum, cell state specificity in primary hematopoietic progenitors is driven by specific negative synergies between TFs.

### Frequent loss of repressor function in K562 cells

To compare these results to a cell line context, we first focused on the single factor Library A, screened in leukemia-derived K562 cells. At the level of activators, there was good agreement between primary HPCs and K562: 17/22 activators identified in HPCs were also activators in K562 cells, and the fraction of sequences activated by each factor in the two cell types was quantitatively similar (Figure S7a,b: Pearson R: 0.82). Trp53 behaved noticeably different between HPCs and K562 cells, as primary hematopoietic cells activate p53 upon lentiviral infection^47^ without affecting the differentiation of cells in the *ex vivo* system used here (ref. ^27,29^ and Figure S2). By contrast, p53 is mutationally inactivated in K562 cells^48^. At the level of repressors, however, there was little agreement between HPCs and K562. Gfi1b, Gfi1, Runx1, Meis1, Meis2, Pbx1 and Tcf12 all lost their repressive ability in K562 cells (Figure S7a,c). Factors associated with occupancy dependent duality (Elk1, Sp1, Creb1) behaved very similar in K562 and primary HPCs (Figure S7d).

Together, these results suggest that the function of several repressors is impaired in K562 cells.

### Neutralizing interactions are widespread in primary HPCs, but not K562s

To investigate how these alterations affect negative synergies between TFs, and to evaluate how widespread negative synergies between factors are across a broader set of TFs, we turned to Library C, where all pairwise combinations of 42 TFs were covered. Library C had a large complexity and lower number of sequences per TF pair, and was primarily analyzed aggregated across HPC cell states (Figure S8a,b). The estimates of activation strength per factor combination were highly concordant between Library B and C (Figure S8c, Pearson R: 0.87).

For each factor pair, we computed the ability to induce repression or activation beyond the thresholds defined by random DNA in both HPCs and K562 (Figure 7a,b, Table S3). This revealed that more combinations resulted in inactivity in HPCs, compared to K562 (Figure 7a-c). Specifically, combinations of activators and repressors were usually inactive in HPCs, but often displayed activity in K562 (Figure 7d), also when controlling for different levels of measurement noise between primary cells and K562 (Figure S8d,e). Only Trp53 was able to activate transcription in virtually all combinations in HPCs, whereas Elk1, Fli1, Nfyc, Klf1/Klf4 and Sp1 were always activators in K562. These global analyses suggest that TF interactions are frequently neutralizing in primary cells, but not in a leukemia cell line.

**Figure 7:**
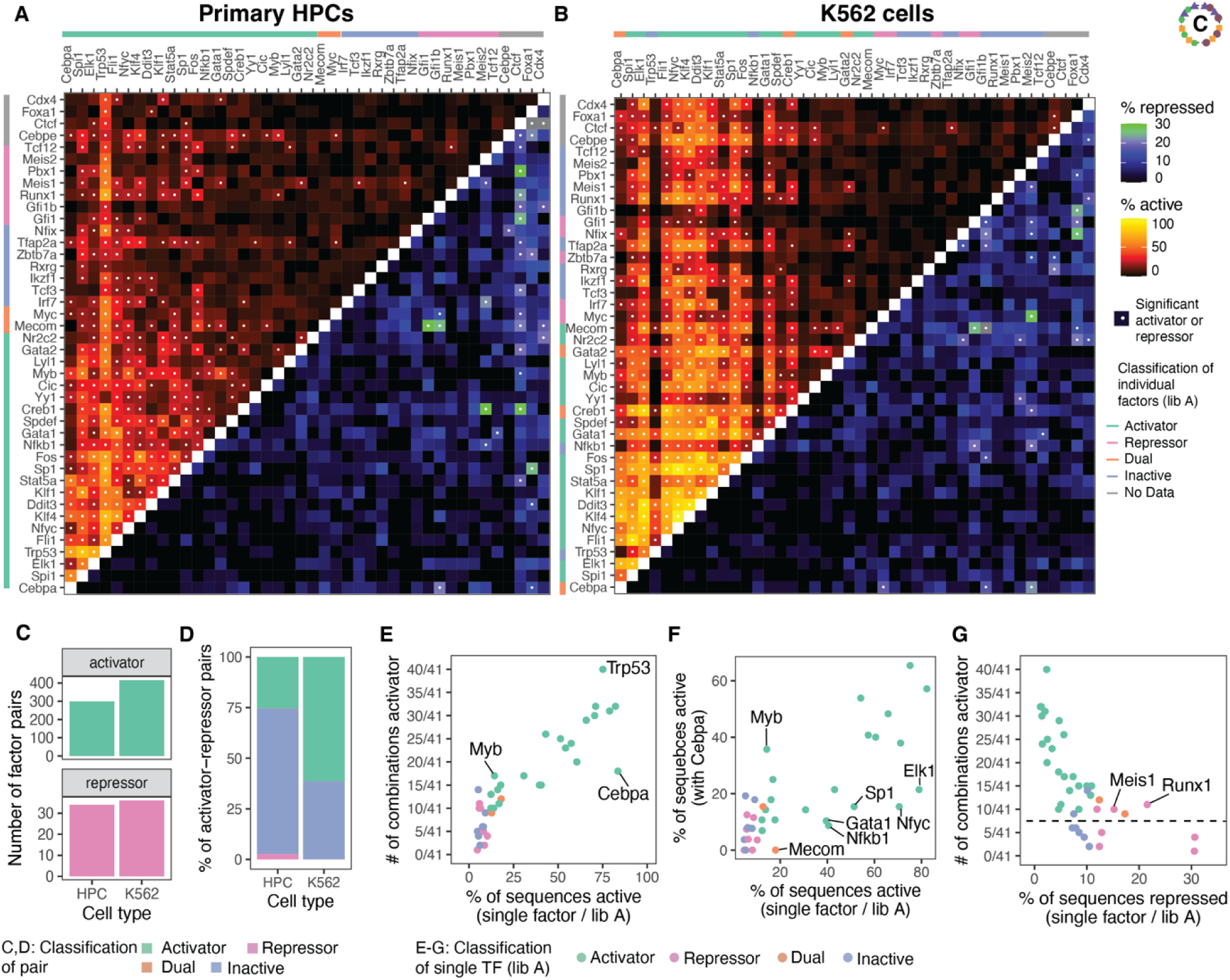
Neutralizing interactions between TFs are widespread in primary hematopoietic progenitors, but lost in K562 cells. See Table S3 for source data. Analyses are based on Library C, see Figure 1d for design. **A.** Heatmap indicating, for each factor pair, the maximum fraction of sequences active or repressed. Dots indicate that a factor combination was classified as activator (upper triangular matrix) or repressor (lower triangular matrix). **B.** Same as panel A, except that data from K562 cells is shown. **C.** Bar chart depicting the number of factor pairs classified as activator or repressor in the two cell types. **D.** Bar chart depicting the functional classification of activator-repressor pairs in the two cell types. **E.** Cebpa is frequently inactivated by other factors. Scatter plot depicting the relationship between the activity elicited by each single factor in the context of random DNA (x axis, see also Figure 2a) and the number of TF combinations in which the factor acts as an activator (y axis). **F.** Scatter plot depicting the relationship between the activity elicited by a single factor in the context of random DNA and the activity elicited by the factor in combination with Cebpa (y axis). **G.** Runx1 and Meis1 are repressors that support activation in some combinations. Scatter plot depicting the relationship between the repression elicited by each single factor in the context of random DNA (x axis) and the number of TF combinations in which the factor acts as an activator (y axis). Dashed line serves to guide the eye.

To characterize these interactions between motifs as unspecific/additive or specific/synergistic, we exploited the observation that generally, the ability of a factor to activate transcription in combination with other binding sites was well predicted by its ability to activate transcription on its own (Figure 7e, S8f). Stronger activators were active in more combinations, compared to weaker activators. In HPCs, Cebpa strongly defied this relationship and was specifically neutralized by several motifs, including strong activators such as Gata1, Nfyc and Elk1, and it was entirely inactive in combination with Mecom (Figure 7f). These neutralizing interactions were lost in K562 cells. Rather, Gata1 sites increased Cebpa’s activity in that context (Figure S8g). Interestingly, Myb served as a specific co-activator for Cebpa, Gata1 and Nfyc in both cellular contexts (Figure 7f, S8g-i).

To complement this analysis, we also investigated the sensitivity of single factor measurements from Library A to random DNA background (Figure S9, Supplementary Note 2). These analyses independently confirmed that “GATA” occurrences in the background of constructs containing Cebpa binding sites reduced their activity in primary cells but not in K562 (Figure S9f,g).

Together, these analyses highlight examples of specific TF interactions that are dysfunctional in the leukemia setting.

Runx1 and Meis1 were repressors on their own that were permissive of activity in a larger number of combinations, compared to other repressors (Figure 7a,g). Interestingly, these factors activated transcription in combination with weak activators (Gata2, Myb), factors that on their own were inactive (Tcf3, Tfap2a), or even repressors (Runx1-Tcf12 combination), indicating that they can turn from repressors into co-activators depending on context (Figure 7a, S8j,k). By contrast, Runx1 and Meis1 elicited weak or no transcriptional activity in combination with strong activators such as Cebpa, Elk1, Nfyc and others. Again, these specific interactions were lost in K562, where Runx1 and Meis1 sites were permissive of activation by strong activators, and inactive in the context of other factors. Finally, Ctcf acted as a strong repressor in HPC. Ctcf is known to recruit co-repressors^49^, but lost this ability in K562s.

Together, these analyses suggest that in primary HPCs, transcription factors frequently neutralize each other’s effect, and only specific combinations of factors are permissive of expression. By contrast, in K562 cells many combinations of factors were able to drive transcription.

## Discussion

During hematopoiesis, stem cells repress differentiation associated genetic programs whereas progenitors express these programs in a highly stage-specific manner. By contrast, in leukemia stem- and progenitor programs are co-expressed. It appears paradoxical that this specificity and functional duality can be achieved by master TFs that are expressed in a highly overlapping manner throughout the, healthy and diseased, differentiation landscape. Here, we have constructed enhancers from scratch to identify three principles enabling the design of cell state specific enhancers:

First, motifs of a single TF can act as activating or repressive, simply depending on the (predicted) occupancy of the TF on the enhancer. This *occupancy dependent duality* enables enhancers to act as band pass filters for TF activity. Non-monotonic responses of reporters to increased numbers of TF sites are sometimes phenomenologically described as negative homotypic interactions^17,20^, and sometimes referred to as ‘squelching’, which originally described the sequestration of co-factors caused by the over-expression of a transcription factor^50,51^ and is not an appropriate term to use here. The systematic design of our large-scale enhancer libraries, which explore a wide range of affinities and binding site numbers of TFs, allowed us to instead suggest differential modulation of distinct biochemical steps in a polymerase activation cycle as a minimal plausible mechanism of occupancy dependent duality. This model is also supported by recent data on differential TF-cofactor dependencies^52^.

Second, motifs of a single factor can be interpreted differently in different cell states. Such *cell context dependent duality* can emerge from occupancy dependent duality, as observed in the case of Myc, where a low TF dose is required for activity. However, cell state dependent duality can potentially also emerge from a different co-factor environment in different cell states, or differences in splicing (described for Mecom^53^) or post-translational modifications.

Third and most importantly, motifs of two different factors can interact non-additively. While the effects of TFs as activators or repressors are well-known to be context dependent^54^, this is, to our knowledge, the first demonstration that combinations of activators alone can give rise to a repressor. Previous work used fusions of individual TF effector domains to a DNA binding domain to demonstrate widespread neutralizing interactions between activator domains, and dominance of repressor domains over activators^55^. Here, we demonstrate that these principles can result in TFs that have no apparent autonomous repressive ability but become repressors in combination. Such *combinatorial duality* is a powerful principle enabling enhancers to sense TF ratios and thereby become cell state specific, even if constituent TFs are expressed and active throughout differentiation. Interestingly, both adult^2,4^ and embryonic^5^ stem cells are characterized by a co-expression of lineage factors. When these combinations result in repressors, a repression of differentiation programs as well as the mutual exclusivity of stem- and progenitor programs is ensured.

This design principle has previously been overlooked, possibly because functional genomic studies in the context of gene regulation are often performed in leukemia or cancer cell lines. Leukemia is frequently driven by mutations in epigenetic modifiers or signaling master regulators^56,57^ that lead to gross alterations of the epigenetic landscape^58^ and an epigenetic environment widely permissive of activation. By contrast, primary blood progenitors have been described to have two orders of magnitude fewer co-activator protein molecules in the nucleus, compared to co-repressors^59^. Our data suggest that these alterations cause a difference in TF function: Interactions that resulted in inactivity or repression in primary cells were activating in leukemia-derived K562 cells. Such epigenetic alterations can potentially explain the co-expression of stem- and progenitor genes^11^ frequently observed in leukemia.

In sum, our work introduces principles and rules underlying the function of cell-state specific hematopoietic enhancers. Unlike recent studies that used deep learning for designing cell-type specific enhancers in the fruit fly^60,61^, the design rules identified here are fully transparent. Synthetic enhancers characterized here have a well-defined motif composition and can establish specificity from highly overlapping TF expression patterns.

## Data availability

Raw and processed MPRA data are available at https://doi.org/10.6084/m9.figshare.25713519.v1 Data can be interactively browsed at http://veltenlab.crg.eu/MPRA

## Code availability

Scripts used for processing of MPRA data are available at https://github.com/veltenlab/MPRAscripts

## Acknowledgements

We thank Bas van Steensel, Sergi Beneyto, Max Trauernicht and Jessica Velten for providing feedback and discussion throughout the project, and Arnau Sebe, Alejo Rodriguez-Fraticelli, Joseph Bowness, as well as all members of the Velten lab for discussion of the manuscript. We thank the CRG facilities for Genomics and Flow Cytometry, as well as the PRBB animal house, for experimental support. This study was financed by grants from the Spanish National Agency for Research (Grant PID2019-108082GA-I00 to L.V.) and the European Research Council (ERC-StG AI4SYN to L.V.). The authors acknowledge support of the Spanish Ministry of Science and Innovation to the EMBL partnership, the Centro de Excelencia Severo Ochoa and the CERCA Programme / Generalitat de Catalunya.

## Author contributions

RF and LV conceived of the study. RF, ABM, CST and JR performed experiments. RF and LV analyzed the data, with contributions from JR. FPP performed deep learning-based analysis. RMC performed biophysical modelling. RF and LV wrote the manuscript, with contributions from all authors.

## Methods

### Selection of transcription factors and their binding sites

Transcription factors with variable activity and expression during hematopoiesis were selected based on differential gene expression (DGE) using single-cell RNA-seq data from ref. ^27^, as well as differential chromatin accessibility using single cell ATAC-seq data ref. ^62^ (Figure S1a,b). From these analyses, as well as review of the relevant literature^14,35–42^, we selected 42 TFs based on their differential expression/accessibility in single and multiple cellular lineages. For each of these factors we selected the all position weight matrices (PWM) available in Hocomoco v11^63^, Jaspar2020^64^, as well as cis-bp v2.0^65^. We then used the TFBStools^66^ and universalmotif^67^ packages to select the centroid motif for each TF. Next, we compared each of the 42 motifs to each other by computing Kullback-Leibler divergence. The original PWM is depicted for each individual factor with ggseqlogo^68^ (Figure S1c). We also visualized the results with complexheatmap^69^ to identify clusters of similar motifs (Figure S1d).

### Design of DNA libraries

After selecting the key transcription factors from gene expression and chromatin accessibility data of the hematopoietic system, we constructed three synthetic enhancer libraries. We focused on examining the contribution of design principles, like orientation, spacing and number of repeats for each individual TF binding site using synthetic DNA sequences. To not introduce a systematic bias into our GRE library, we decided to use random background DNA. The reason for this is that cryptic TF motifs in a fixed background sequence might have synergistic or antagonistic effects to only a subset of TFs, ultimately leading to biases. We designed homotypic Library A, focused on individual TFs, and heterotypic Library B and Library C, focused on pairs of TFs.

For all libraries, three different components were needed for generating a single sequence, the TF binding sites, spacers between binding sites, and DNA to fill up to a common length. To generate TF binding sites of different affinities, we drew 100.000 sequence motifs from a given PWM and then ranked them by their likelihood given the original PWM (Figure S1e,f). We then sampled sequences in a ± 2.5 % percentile range surrounding a target affinity percentile, where the 100^th^ percentile corresponds to the pest possible motif. This sampling generates different motif instances with a similar affinity.

To generate spacers between binding sites, we generated a list entry with up to 10.000 random DNA sequences for each spacer length used. Spacer sequences larger than 10 base pairs were additional filtered for the occurrence of motif instances of all 38 selected TFs using Fimo^70^. Transcription factors binding sites (TFBSs) and spacers were then assembled from the 3’ of each GRE by alternating TFBSs and spacers. We started with a motif instance for each GRE and due to this the last TFBS in our construct always had the same distance to the promoter.

Finally we generated a filling sequence, so that all GRE sequences have the same length (232, 216 and 220 base pairs for Library A, B and C respectively). Like the spacers, filling sequences were created by generating random DNA and filtering for the occurrence of motif instances of the selected TFs. For each enhancer we generated, we created 100 candidate sequences *in silico* and sampled from the sequences with the smallest number motif occurrences of other TFs. As a last step we added the 15 base pair adaptor sequences from lentiMPRA^31^ for PCR and amplification to each GRE in the 5’ and 3’.

Based on this common design strategy, we created three libraries (see also Table S1):

Library A investigated 38 TFs and for each of them we generated 270 sequences, divided into 3 spacer, 3 orientations, 6 affinities, 6 repeat numbers groups. In total we placed between 1 and 6 binding sites with spacer lengths of 4, 10 and 20 base pairs, to have closer and larger interaction spaces. We sampled the TF affinity from the 10^th^, 25^th^, 50^th^, 75^th^, 90^th^ and 100^th^ percentile (100^th^ quantile being the highest likelihood). For orientation we used only forward and only reverse orientation for all number of repeats, but for 4 to 6 repeats we also designed a tandem arrangement of forward-reverse for the TFBS.

Library B investigated 10 TFs and we created 45 pairwise combinations between these TFs (“TF1 – TF2”). The design space was the following:

a. Containing one or three binding sites for each TF
b. Consisting of combinations of medium (40^th^ - 70^th^ percentile) and high (80^th^ - 100^th^ percentile) affinity binding sites
c. For combinations of three binding sites per TF we explored short spacings (2, 4, 6, 8, 10, 12, 14, 16 and 18 base pairs) and for combinations of single binding sites we explored additional spacings of 32, 64 and 128 base pairs, see Table S1
d. All combinations of forward and reverse orientations for both TFs
e. For constructs with 3 + 3 binding sites, the arrangement can be either alternating between TF1 and TF2 or all sites for a TF a placed together
f. Placing TF1 at the 3’ end of the construct, or placing TF2 at the 3’ end

For combinations involving Heptad factors (Fli1, Gata1/Tal1, Gata2, Lyl1, Meis1, Runx1), or Spi1, we generated 912 sequences and combinations involving also Trp53, Cebpa or Gfi1b we created 336, by exploring more or less options at step b and e, see Table S1.

Library C profiles 861 combinations of 42 TFS. It has identical design parameters as Library B, but the spacing range for single binding factors Library B (step c with 32, 64 and 128 base pairs) was extended to 2-200 base pairs. For this library we sampled from the whole design space to create 22-34 sequences per TF pair.

Library B and Library C also contain a section of additional control sequences.

a. For some of the generated sequences we generated permuted versions. We shuffle the whole sequence, just the TFBS, the background (everything besides the TFBS), or the filling sequence for 200 randomly selected sequences
b. For 1,500 randomly selected sequences in Library C, we also generated reverse complement sequences
c. We always included the Trp53 sequences from Library A
d. We included inert, as well as a positive control sequence^71^

In summary, we created synthetic GRE sequences for 38 key hematopoietic transcription factors. To get a detailed understanding of each of these factors we investigate them on their own (Library A) as well as in pairs (Library B and Library C).

### Cloning of DNA libraries and GRE-Barcode association libraries

DNA libraries were ordered from Twist Bioscience and cloned into a lentivirus reporter plasmid that contains a I-SceI site and a EGFP reporter gene, pLS-SceI (Addgene, Plasmid #137725) as described in ref. ^31^, with some adaptations. Briefly, an initial qPCR was carried out to determine the number of cycles that is required to amplify the oligo pool in the first-round PCR, where vector overhang, minimal promoter, and adaptor sequences are added. We extended the denaturation step during cycling from 15 seconds to 20 seconds and 12-17 cycles were used for each of our DNA libraries. Once a cycle number was determined, additional PCR reactions were performed to generate at least 100 ng of the first-round PCR product. In the second-round PCR, 4 or 6 cycles were used and multiple PCR reactions were set up to generate at least 250 ng of product, where a 15-bp barcode and a vector overhang sequence downstream of the first-round PCR fragment were added. Similarly, we extended the denaturation step during cycling from 15 seconds to 20 seconds. The MPRA Library A used for HPCs underwent an additional PCR prior the first PCR using 20 cycles with the same conditions. Following library amplification, vector was linearized with AgeI-HF and Sbfl-HF and the undigested vector was eliminated by I-SceI digestion. The linearized vector and the purified insert DNA were then recombined. Typically, 100 ng of the assembled plasmid was electroporated into 100 µl of NEB10-beta electrocompetent cells using the Gene Pulser Xcell System (Bio-Rad). Pre-warmed Stable Outgrowth Medium was added to the electroporated cells and topped up to 4 mL in total. After an one-hour recovery at 37C, 500 µl of bacteria was plated onto a single 25cmX25cm LB agar plate with 100 mg/mL ampicillin for a total of 8 plates. To verify the plasmid sequence, individual colonies were picked, purified with the QIAprep Spin Miniprep Kit (Qiagen, 27106), and Sanger sequenced. Colonies were scraped off from each plate and collected for purification with the PureLink HiPure Plasmid Filter Maxiprep Kit (Invitrogen). Finally, to associate the barcode to the GRE, a PCR reaction was set up on the plasmid library to add sequencing adaptors and samples indexes to the GRE-barcode fragment. This PCR product is then gel-purified and submitted for pair-end sequencing to determine a dictionary of GRE-Barcode associations (see section *Data processing: GRE-Barcode association libraries*).

### Virus production

Lentivirus was produced in HEK293FT cells combining library plasmids (1.64 pM), psPAX2 (1.3 pM) and pMD2.G (0.72 pM) (Addgene plasmids #12260 and #12259). 6h transfection was performed with Lipofectamine 3000 following the protocol of the manufacturer. Lentivirus were collected 72h after transfection, filtered with low protein binding 0.45 µm filter and precipitated with sucrose cushion ultracentrifugation^72^ (60000 g for 1.5 h at 4°C). Lentivirus was resuspended in StemSpan media, aliquoted and stored at -70°C. All lentivirus were titrated with Lenti-X qRT-PCR Titration Kit (Takara Bio 631235). Before MPRA screens, all lentivirus preparations were tested in K562 cells and HPCs to assure lentiviral efficiency, cell viability and activity of the GFP reporter.

### Extraction, culture and infection of primary HSPCs

For each MPRA screen, 5 male and 5 female C57BL/6 mice (Charles River Laboratories) at 3 months of age were euthanized by cervical dislocation. Hematopoietic stem and progenitor cells were freshly isolated from bone marrow. First, long limb bones were isolated and cleaned from connective and muscle tissue. During the entire manipulation, bones and cells were kept in 2% FBS-PBS on ice. Bones were crushed and washed with 2%FBS-PBS until bones were completely white. Collected cells were filtered through 40 µm strainer and centrifuge at 400 g 5 min at 4°C. Red blood cells were lysed with ACK Lysing Buffer (Gibco) at a ratio of 1 ml/mice. Lineage negative (Lineage cell depletion kit, Miltenyi Biotec) and cKIT positive (Cd117 Microbeads mouse, Miltenyi Biotec) cells were isolated from the total bone marrow cells by two steps of magnetic cell sorting (Miltenyi Biotec). HSPCs were cultured for 4 days in Ultralow attachment plates (Corning) at 37°C and 5% CO2 conditions. HSPC media was adapted from ref. ^27^ as follows: StemSpan media (StemCell Technologies) supplemented with 1% penicillin-streptomycin and the following cytokines: mSCF (50 ng/ml) (Peprotech), hEPO (60 ng/ml) (Peprotech), Il-11 (50 ng/ml) (Peprotech), FLT3 (50 ng/ml) (Peprotech), TPO (50 ng/ml) (Peprotech), Il-3 (20 ng/ml) (Peprotech), Il-5 (10 ng/ml) (Peprotech) and Il-6 (10 ng/ml) (R&D Systems). 100 million HSPCs were infected 24h after isolation with Library A (1.09*10^11^ mRNA lentiviral particles), Library B (3.3*10^12^ mRNA lentiviral particles) and Library C (1.24*10^12^ mRNA lentiviral particles) for 20h, afterwards cells were washed and allowed to recover for 48h.

### Culture and infection of K562 cells

K562 cells were cultured in RPMI media supplemented with 10% FBS and 1% Penicillin-Streptomycin. For K562 cells screen, 2 million cells were infected with each Library A, Library B and Library C (the amount of mRNA lentiviral particles ranged between 2.2*10^9^ and 1.63*10^10^ for all libraries). This procedure was done in duplicates. Cells and virus were co-incubated for 20h, then cells were washed and let them recover for 24h. K562 cells were collected after 2 days of transduction. Before nucleic acid isolation, cells were washed twice with PBS.

### CITE-seq and gating scheme design

To design a gating scheme for the *ex vivo* HPC culture system, we used a data-driven strategy we had developed earlier^73^. In brief, a single cell CITE-seq reference map of HPC differentiation cultures was created, linking the expression of surface markers to the cellular identity. Hereby, we use the single cell RNA-seq data to define cellular states based on their gene expression profile and designed a surface marker panel to isolate cellular states of interest. We included 26 cell type and differentiation specific antibodies and performed three Cite-Seq experiments using the 10X Genomics platform, specifically the single cell 5’ kit together with Biolegends TotalSeq-C antibodies (see Table S4). Data were analyzed as described^73^. We annotated the cells based on mapping it to a reference^74,75^ (Figure S2a), as well as annotating the cells manually (Figure S2b), based on marker expression (Figure S2d) and antibody expression (Figure S2e) across myeloid lineages. Our next aim was to generate an antibody gating scheme to sort the hematopoietic culture system into distinct cell stages. For this we aggregated the cells into seven clusters, based on their gene expression profile. We then used to antigen expression matrix to construct regression trees, as described^73^, thereby defining a FACS gating scheme. We then refined this scheme based on experimental considerations and flow cytometry experiments, to obtain the scheme shown in Fig S2f,g. The gating scheme was validated using FACS sorting and a modified smart-seq2 protocol^76,77^ (Figure S2h. A more detailed description of the CITE-seq based characterization of the differentiation cultures is provided in Supplementary Note 1.

### Cell sorting

HPCs were stained to identify and sort seven cell states as detailed in Figure S2f. For the staining, the following conjugated antibodies were used: Cd55-AF647, cKit-Bv605, Cd274-PE, Itga1-APC/Fire 750 and Ly6a-Bv785 (Table S4). Additionally, DAPI was used as a viability dye. A minimum of 2 million live cells for each population were sorted using FACSAria II cell sorter (Becton Dickinson, San Jose CA). Two replicates were sorted for each population.

### Bulk RNA-seq

To assess the reproducibility of our culture conditions in each of our screens, we performed bulk RNA-seq by taking a small aliquot of the total RNA from the MPRA library preparation from sorted populations and generating full-length transcriptomes with a modified smart-seq2 protocol^76,77^. Briefly, 10 ng of total RNA was lysed in lysis buffer without Triton X-100 followed by reverse transcription and PCR preamplification with 12 cycles and only the Smart-seq2 ISPCR primer. The amplified cDNA was then purified using CleanPCR beads (CleanNA, CPCR-0050) and tagmented with a homemade Tn5^78^. Lastly, PCR was performed to add indexed sequencing adaptor sequences to the final library. Sequencing data were analyzed using kallisto^79^.

### MPRA library preparation

We followed the lentiMPRA protocol^31^. In brief, sorted HPC and K562 cells were lysed with RLT-1% β mercapthoethanol lysis reagent for DNA and RNA isolation. To increase the efficiency of cell lysis, 700 µl of RLT-1% β mercapthoethanol were added to each sample, the lysates were homogenized using a syringe and 21-gauge needle and stored at -70°C for later processing. RNA and DNA were isolated following the manufacturer’s protocol (Allprep DNARNA kit, Qiagen). For RNA samples, we included a DNAse treatment between the washing steps. Once eluted, for RNA samples, we performed a second rigorous DNAseI treatment following manufacturer’s protocol (Turbo DNA-free kit, Qiagen). RNA was converted into DNA by reverse transcription. At this step, 16-bp UMI and P7 flowcell sequence were added. First, RNA was incubated with P7-pLSmP-ass16UMI-gfp and dNTPs for 5 min at 65°C. RT reaction was performed with SuperScript IV at 55°C for 10 min followed by an inactivation step at 80°C for 10 min. To prepare and amplify the library for sequencing, we did a 3 cycles PCR in which P5 flowcell and indexing sequences and 16-bp UMI and P7 flowcell sequence were added, independent indexes were used for each replicate and population (see Table S5 for primer sequences). We determined an optimal cycle number (13-20 cycles) by qPCR to amplify and added P5 and P7 sequences in a second round-PCR. Before sequencing, all samples were bioanalyzed to check the size and the purity of the product.

### Data processing: GRE-Barcode association libraries

Data from GRE-Barcode association sequencing contained a forward and reverse read covering the GRE, and an index read covering the barcode in the 5’end of the GFP transgene (see Figure 1b). Forward and reverse reads were mapped against the synthetic GRE sequences using bwa mem^80^. The resulting .bam file was then processed with a custom Perl script to create a GRE-Barcode dictionary (see code availability statement). We first counted the reads supporting each Barcode-GRE association. For barcodes assigned to multiple GREs, we sorted GREs by the number of reads, selected the most supported assignment, and counted the number of reads supporting deviant GRE assignments for subsequent filtering. We also determined the mean alignment quality for each Barcode-GRE association to identify clones with sequence errors in the GRE. Next, we corrected barcodes for sequencing errors. To that end, we sorted all barcodes associated with a given GRE by the number of reads supporting that assignment, and removed barcodes with a hamming distance of one to a more strongly supported barcode. This processing step resulted in a CSV file including the (corrected) barcode, GRE name, number of reads supporting the assignment, number of reads supporting a deviant assignment, as well as minimal, mean and maximal alignment score. We then filtered this dictionary by alignment scores (between 290 and 292), as well as the number of reads supporting an assignment (has to be five times larger than the deviant reads supporting other assignments). Barcodes containing homo-nucleotide stretches, more than 10 identical nucleotides per barcode, were additional filtered out. Filtering criteria were empirically evaluated, see below (section *Empirical evaluation of MPRA processing parameters*).

### Data processing: MPRA libraries

MPRA barcode sequencing data were processed with a custom Perl script (see code availability statement). In brief, these data contained a forward and reverse read of the GRE barcode in the 5’UTR of the GFP transgene, as well as an index read containing a unique molecular identifier (UMI). Our script first loads the filtered GRE-Barcode association dictionary. We then iterate through all MPRA sequencing reads and verified that the forward and reverse barcode read are identical and match any assigned barcode perfectly. Read counts of all UMIs observed in combination with any given barcode are stored. After processing all reads, UMIs associated with a specific barcode are sorted by supporting read count and error correction is applied as in the preceding section. This processing step resulted in a CSV file including GRE name, number of UMI, number of reads, as well as additional columns for number of UMIs covered with different read thresholds. Similar CSV files were generated both for libraries created from RNA, and libraries created from DNA.

### Empirical evaluation of MPRA processing parameters

Both the GRE-BC association dictionary, as well as the MPRA Barcode counting table, can be subject to different quality filtering and processing options, which need to be adjusted to sequencing depth, infection rate, cell number, and other technical parameters, and are usually set arbitrarily. We here exploited the fact that MPRA activity measurements in our synthetic libraries are predictable from sequence design to instead choose optimal parameters systematically.

To that end, we performed a parameter grid search for each MPRA experiment. The grid search is performed to find the processing parameters that result i) in the highest R^2^ between replicates, ii) the highest R^2^ of a random forest model predicting activity from design and iii) the highest number of GREs passing filter. We explored the following processing parameters:

1. Different thresholds for the minimal number of reads in the GRE-BC association
2. Different minimum number of reads per UMI
3. Whether UMI counts per BC on DNA should be capped at 1, reflecting the assumption that at the complexity of barcodes used here, a single barcode is likely to reflect a single integration site
4. Whether a BC that is observed in RNA, but not in DNA, should be counted as observed in DNA with one UMI
5. Different minimum number of UMIs per GRE (on DNA and RNA)
6. Normalization of RNA barcode counts to the DNA barcode counts from the MPRA experiment, versus normalization to the read count from the GRE-BC association library (as typically done in STARR-seq^81^)

Table S6 details the final set of parameters used.

For each GRE in a respective cell state we will compute a library size normalized count value for DNA/RNA for both replicates. We refer to the final log2 ratio between library size normalized RNA and DNA counts as the raw activity measurement for a given GRE.

### Identification of active and repressed sequences

For the qualitative identification of sequences and factors as repressors or activators, we determined if each sequence was significantly more or less active than random DNA. To this end, for each sample and experiment, we computed the distribution of measured activities for random DNA. For each candidate enhancer sequence, we then determined if the measured activity was higher than the 95^th^ percentile of that distribution, or lower than the 5^th^ percentile (Figure 2a). For a given group of sequences, this allowed us to determine what percentages of sequences was ‘active’ or ‘repressed’ in a given cell state. To summarize this quantity into a single number per factor, we computed its maximum across cell states. We then repeated the same analysis on 5,000 sets of re-sampled random DNA measurements sampled to identical size, and determined a 95% confidence interval. Only factors/factor pairs with a higher frequency of active or repressed sequences, compared to this baseline, were defined as ‘activator’ or ‘repressor’.

### Scaling of the data

We sought to define a robust scale of activity that was quantitatively comparable between different samples and experiments. This was important to account for technical variabilities in cell numbers, infection rates, and library preparation that affect raw MPRA activity measurements. To that end, we exploited the observation that all primary hematopoietic cells activate Trp53 upon lentiviral infection^47^ without affecting the differentiation of cells in the *ex vivo* system used here (ref. ^27,29^ and Supplementary Note 1). We included an identical set of 270 sequences containing Trp53 binding sites into all libraries. These sequences were characterized by a broad range of activities, and a high correlation across different cell types (Pearson R>0.95) and experiments (Pearson R>0.86). We used these measurements to transform all data to a common scale, where 0 is the median activity of random DNA and 1 is the maximal activity induced by Trp53. To that end, we divided all background-subtracted activity measurements by the 95^th^ percentile of the activity of Trp53 sequences. After this adjustment, random DNA controls measured in both experiments displayed a similar mean/variance relationship and a similar dynamic range. This scaling of the data was used for plotting of the data (in Figure 3b, 4 and 6b-d) but did not make a difference in the statistical or machine learning based analyses, which were comparing sequences to random DNA background.

### Affinity computation and data visualization

For each TF binding site placed on random DNA during the library design, we computed the affinity score S from PWM following^82^, using the equation

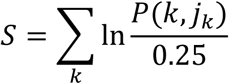

Where j_k_ is the base at position k of the motif and *P*(*k*, *j*_*k*_) is the probability of observing that base at the kth position of the position weight matrix, considering the same PWM as during the design process (see section *Design of DNA libraries*, and Figure S1). Given a PWM, the maximum score *Smax* corresponds to the site sequence that has at each position the most probable nucleotide given the PWM. Then the affinity constant for any other site sequence is a monotonous function of *S* given by^82^:

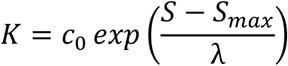

Where *c0* is a constant that gives the correct units. Given the monotonous relationship between *S* and *K*, we refer to *S* as “affinity score”. For the plots in Figure 3b and 4, we then summed this quantity over all the sites placed in the GRE and used smoothened conditional means (geom_smooth function from ggplot2) for plotting. All other plots were created with the ggplot2 or pheatmap packages in R.

### Biophysical models

We used a biophysical model of gene regulation to investigate non-monotonous responses in Creb1, Elk1 and Myc (Fig. 4d-g). In separate work^45^, we extensively characterize this behavior from a theoretical viewpoint. In that work, we show that non-monotonous effects of a TF with respect to binding site affinity can arise when gene regulation is limited at multiple steps, and a TF acts incoherently on those steps, such that it acts on one or more steps to promote transcription, and on other steps to hinder it. In order to see if this same mechanism could account for the non-monotonicity observed here, we used a model where the equilibrium average occupancy of TFs on the regulatory sequence affects the rates of a polymerase cycle model^45^.

To calculate average occupancy, we first calculate the affinity *K_i_* for each site *i* in a sequence, given the TF PWM, as described in the section *Affinity computation and data visualization,* with *c0* and λ as free parameters to be fitted. Then, we consider equilibrium binding, as described^83^. Briefly, we consider the hypercube graph that has as vertices all possible combinations of sites bound (fully unbound sequence, TF bound at site 1, TF bound at site 2,…, TF bound at sites 1 and 2, at sites 1 and 3,…, at sites 1,2,3 and so on), edges between states *u* and *v*, if *v* has one more site bound than *u,* at position *i,* and edge label between *u* and *v* the product of the affinity constant *K_i_* and TF concentration *x*. *x* is taken as another parameter to be fitted. The model assumes independent binding among the sites, although binding cooperativity could be included at the expense of more free parameters. For the fits of the space-dependent Elk1 behavior, we modelled the effects of space phenomenologically, through *x,* so that all affinities would be equally affected by a given spacing. We note that this is just a proxy for space-dependent cooperativities, but avoids the parameter explosion that would involve explicitly accounting for those^83^.

Given any state *s*, we assign it a weight *μ*_s_, given by the product of the affinity constants corresponding to the sites bound in *s*, multiplied by the TF concentration raised to the number of sites bound. We then calculate the average occupancy *a* following^83^ as

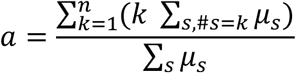

where the inner sum in the numerator runs over the states with k sites occupied, and the sum in the denominator runs over all states.

We then consider a 3-state polymerase cycle where the transition between states 1 and 2 is reversible, and the other irreversible, as in ^84^, so that in the absence of regulation the cycle rates are *k1*, *km1*, *k2*, *k3* and this yields a steady-state transcription rate:

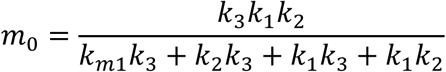

We then assume each rate can be non-linearly modulated by the TF according to:

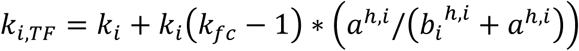

So, for each rate assumed to be modulated (which is decided a priori), there are three free parameters: *k*_*fc*_, *b*, *h*. Note that if *k*_*fc*_ > 1 the TF accelerates that rate, and if *k*_*fc*_ < 1 the TF decelerates the rate. So, we can allow incoherent regulation when the *k*_*fc*_ parameters can all be greater or less than 1, or we can impose coherent regulation by forcing *k*_*fc*_ to be greater than one for the forward rates and less than one for the backward rate. Given the TF-regulated rates, the corresponding TF-regulated transcription rate *m*_*TF*_ can be calculated by using the modulated rates into the previous equation for *m*_0._ We finally assume linear mRNA degradation and calculate the natural logarithm of the fold change in gene expression as *ln*(*m*_*TF*_ /*m*_0_) which is the quantity in the plots and which we compared to the data in order to fit the model.

Fits were performed considering biologically plausible values for *λ* between 0.1 and 10, following^85^ but slightly broadening the ranges, given the lack of accurate measurements for this parameter for our system. We allowed values for the Hill coefficient h to be between exp(- 3) and exp(3), and the rest between exp(-7) and exp(7) to allow for broad but biologically plausible ranges^84^.

In order to fit the model to the data, we used the genetic algorithm in the eaSimple function of the Python package Deap, with the following hyperparameters: population size=500, 100 generations, a standard deviation of 1 for the mutation process, a probability of 0.25 of mutating each parameter and each individual, and a 0.25 probability of crossover. In addition, we run the algorithm starting from 100 different initial conditions and chose the best solution.

### Machine learning based analyses

To characterize the underlying regulatory grammar for Library A and B we constructed tenfold nested, tenfold cross-validated random forests. Our reference model for each library included all design features, resulting in the following equation for Library A:

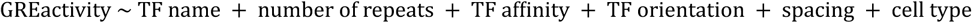

And for Library B:

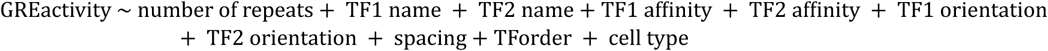

We compared this reference model with ones lacking spacing/orientation information to evaluate the contribution of a given design feature to its resulting activity. Analysis was performed using R packages tidymodels, ranger and parsnip.

We further categorized the hematopoietic TFs of Library A with shape constrained models (R package scam). For each individual TF we fitted the sum of motif affinity and activity with different splines. We compared a flat line with monotonically increasing, monotonically decreasing or concave constraints, using cell type as a covariate. We calculated the Akaike’s information criterion for each fit and categorized each TF with its best fit. We then visualized the delta AIC for each TF to the second-best model (Figure 4b).

## Supplementary Figures

**Figure S1.**
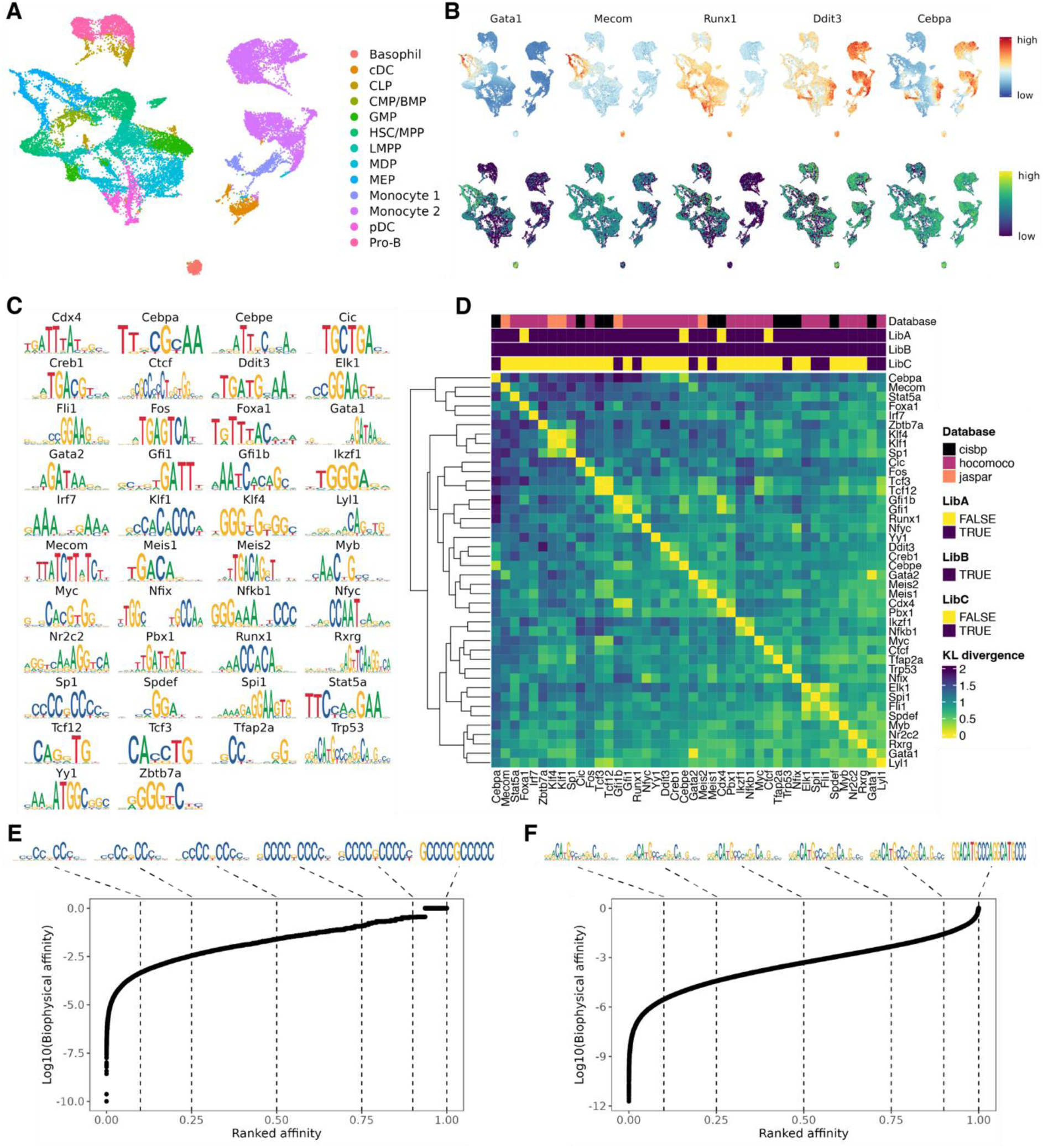
Selection of transcription factors and their binding sites. See also main Figure 1c,d. **A.** uMAP of single cell chromatin accessibility data^62^ of hematopoietic stem and progenitor cells. **B.** Inferred TF accessibility from chromVAR^86^ (top panel), as well inferred gene activity from Cicero^87^ (lower panel) for key TFs Gata1, Mecom, Runx1, Ddit3 and Cebpa. **C.** Selected motifs for the 42 TFs used in this study **D.** Heatmap depicting the motif similarity using the Kullback-Leibler (KL) divergence for the selected motifs. Column annotation highlights the presence in a given GRE library and the row dendrogram depicts the similarity between the motifs. **E.** To generate motifs of a specific affinity, we initially we draw 100,000 sequences for each TF from its PWM and rank them according to their likelihood given the PWM (x-axis). At the same time we can calculate the biophysical affinity (y axis) as described in Method, section *Affinity computation and Data Visualization*. For a given affinity percentile (dashed lines) we then draw from a ± 2.5 % percentile range. The consensus motifs generated by these draws for each affinity percentile is shown on top of each plot. Panel E shows data for Sp1. **F.** Same as panel E, except that data for Trp53 is shown.

**Figure S2.**
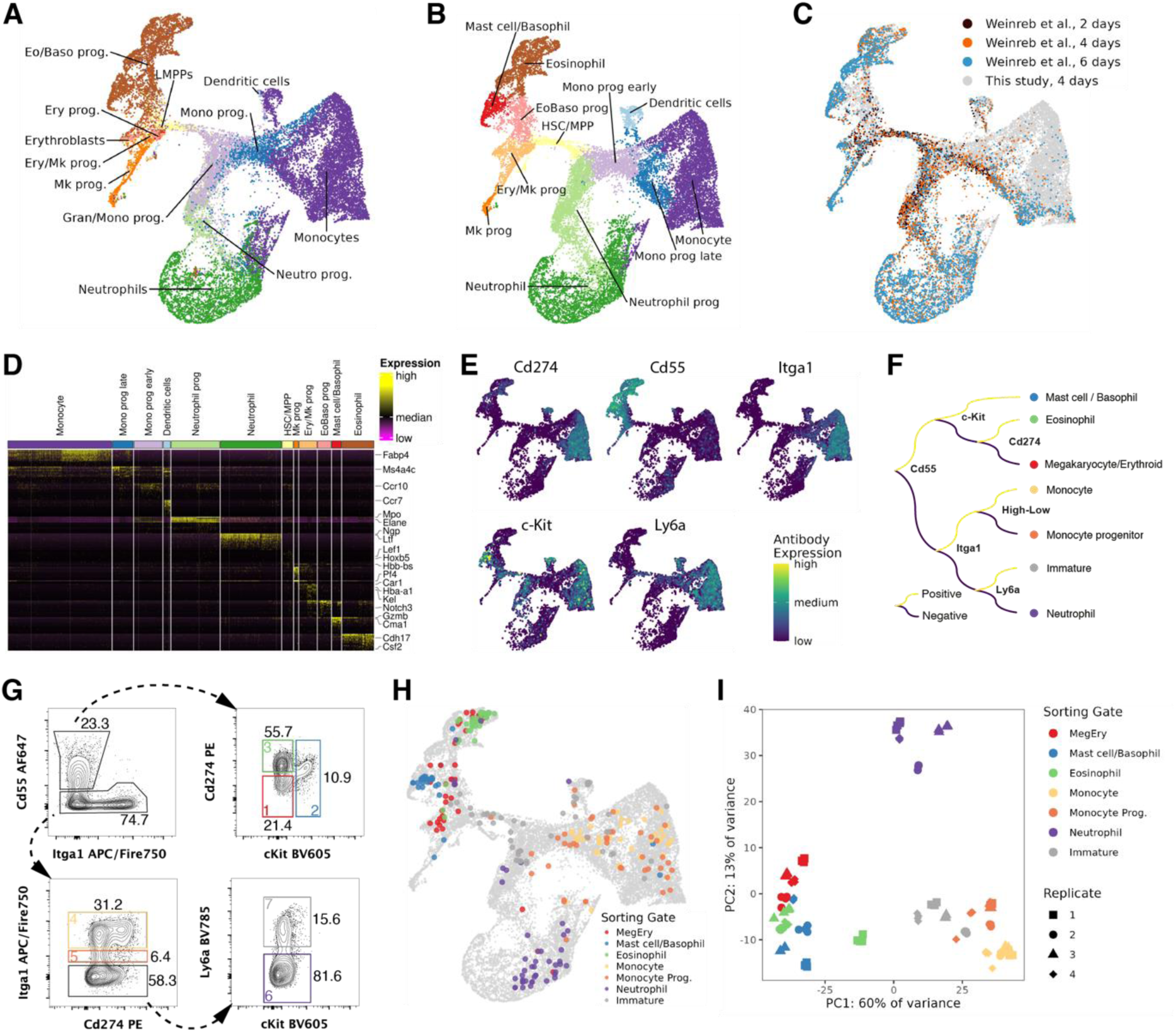
Characterization of cultures and development of a FACS gating scheme. See also Supplementary Note 1, and main Figure 1a,b. **A.** uMAP of single cell RNA-seq data from hematopoietic differentiation cultures. Colors depict label transfer from a reference of primary mouse bone marrow^75^. Prog.: progenitor **B.** Manual cell type annotation of the Cite-Seq data based on marker gene expression (see panel D). **C.** uMAP comparing scRNAseq data of the study that introduced the culture conditions used here^27^, and this study. **D.** Heatmap depicting the expression of highly variable genes, highlighting lineage markers commonly used for the annotation of primary HSPCs. **E.** Display of antibody surface expression. **F.** Data-driven design of an optimal gating scheme using decision trees to divide the culture system into seven major cell types. In total we use five antibodies, Cd55, c-Kit, Cd274, Itga1 and Ly6a. **G.** Implementation of the FACS gating scheme. **H.** Smart-Seq2 validation. We sorted individual cells (based on panel F,G) and performed single cell gene expression measurement of the sorted cells using Smart-Seq2 and remapped the cells back into our reference. **I.** Reproducibility of the cultures. PCA analysis of bulk RNA-seq data collected at different replicate cultures.

**Figure S3.**
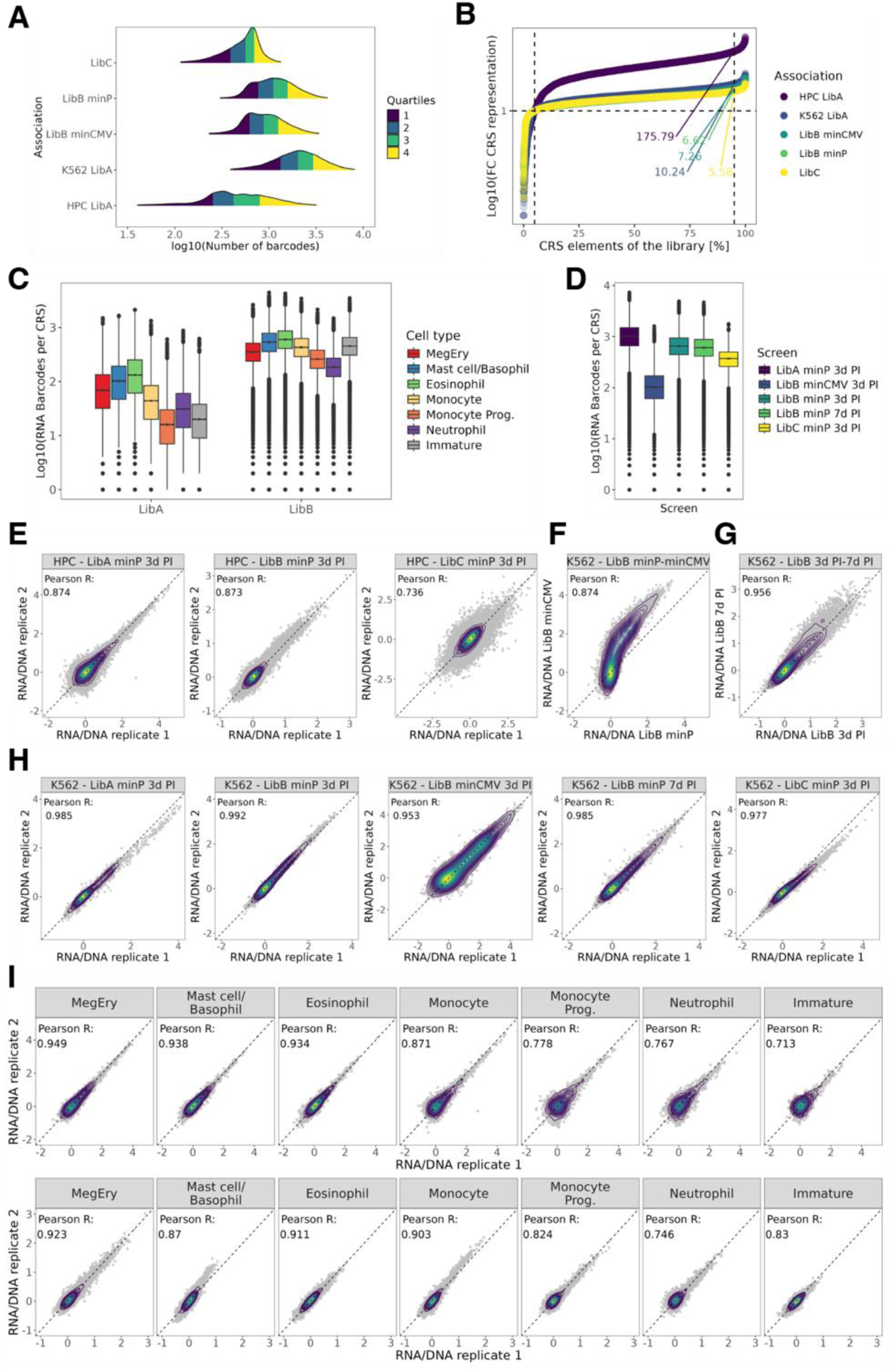
Evaluation of lentiMPRA data quality. See also Supplementary Note 1. **A.** Number of barcodes in the GRE-BC association for each MPRA library. **B.** Evenness of cloning and amplification of the GRE libraries. Only a 5-10 fold difference between the 5^th^ and the 95^th^ percentile was observed, except for Library A in HPCs. **C.** Number of RNA barcodes per element for Library A and Library B in HPCs. **D**. Same as panel C. but characterizing the number of RNA barcodes per element for the five K562 MPRA screens. **E.** Pearson correlation of the technical replicates from the HPC MPRA experiments. **F.** Pearson correlation comparing the GRE activity of different promoters in K562 cells. Enhancer activity of Library B with a minimal promoter (x-axis) compared to Library B with a minimalCMV^88^ promoter (y-axis). **G.** Same as panel F. but comparing the enhancer activity after 3 days post infection (x-axis) with 7 days post infection (y-axis). **H.** Same as panel E. but for K562 cells. Technical replicate correlation of the five MPRA experiments. **I.** Pearson correlation of the technical replicates for each cell type of HSC Library A and Library B. The correlation is between 0.71 and 0.95 for all cell states.

**Figure S4.**
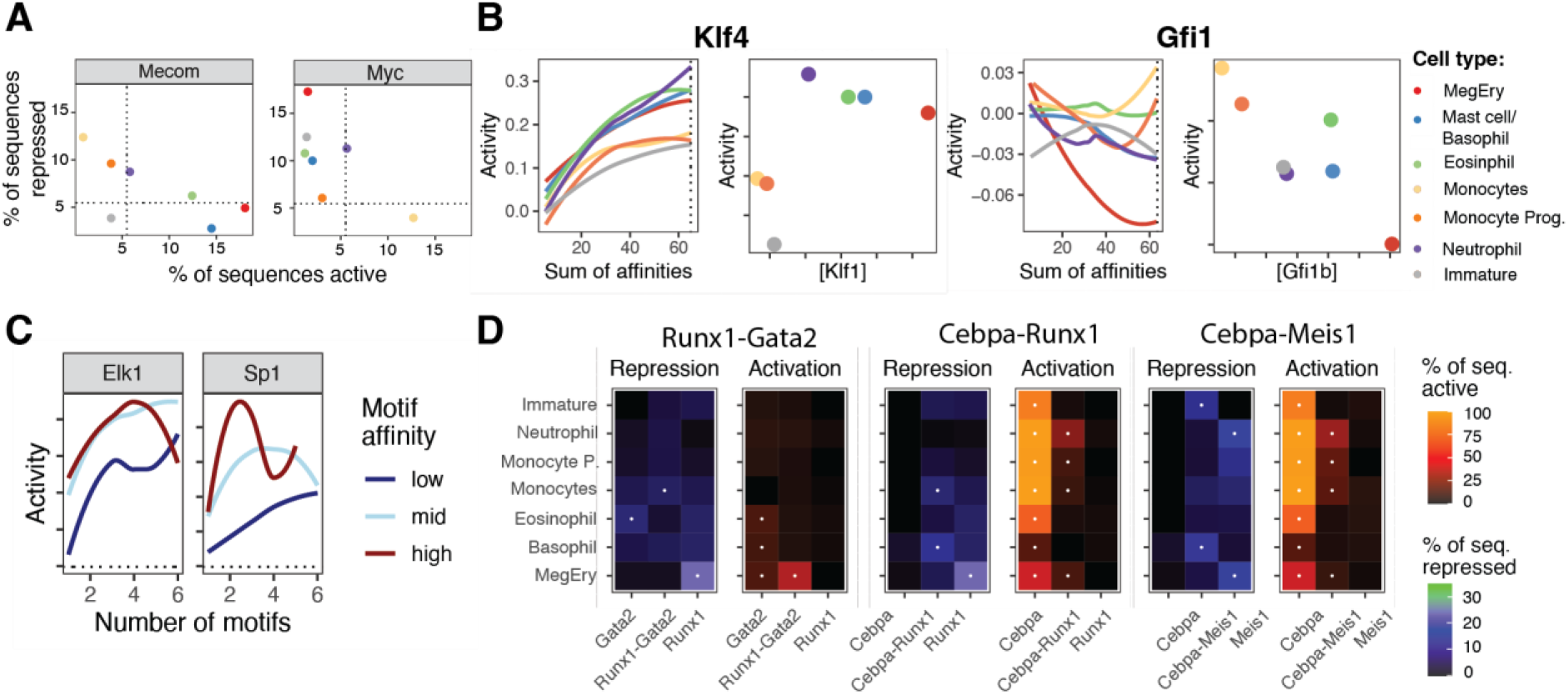
Additional analysis of enhancer activity in Library A and B. See also main Figure 3. **A.** Fraction of sequences active or repressed for constructs containing Mecom and Myc binding sites, stratified by cell type. See panel B for color code. **B.** Left panels: Line plots displaying smoothened means of activity, conditioned by the sum of motif affinities contained in the sequence and stratified by cell state, for Klf4 and Gfi1. See Methods, section *Affinity computation and data visualization.* Right panels: Scatter plots comparing factor expression levels from RNA-seq (x-axis) the activity at the dashed line (y-axis). **C.** Line plots displaying smoothened means of activity, conditioned by the number of motifs and stratified by motif affinity, for Elk1 and Sp1. **D.** Additional examples of activators turning into repressors. For three factor pairs, heatmaps depicting the fraction of active or repressed sequences in the seven cell states.

**Figure S5.**
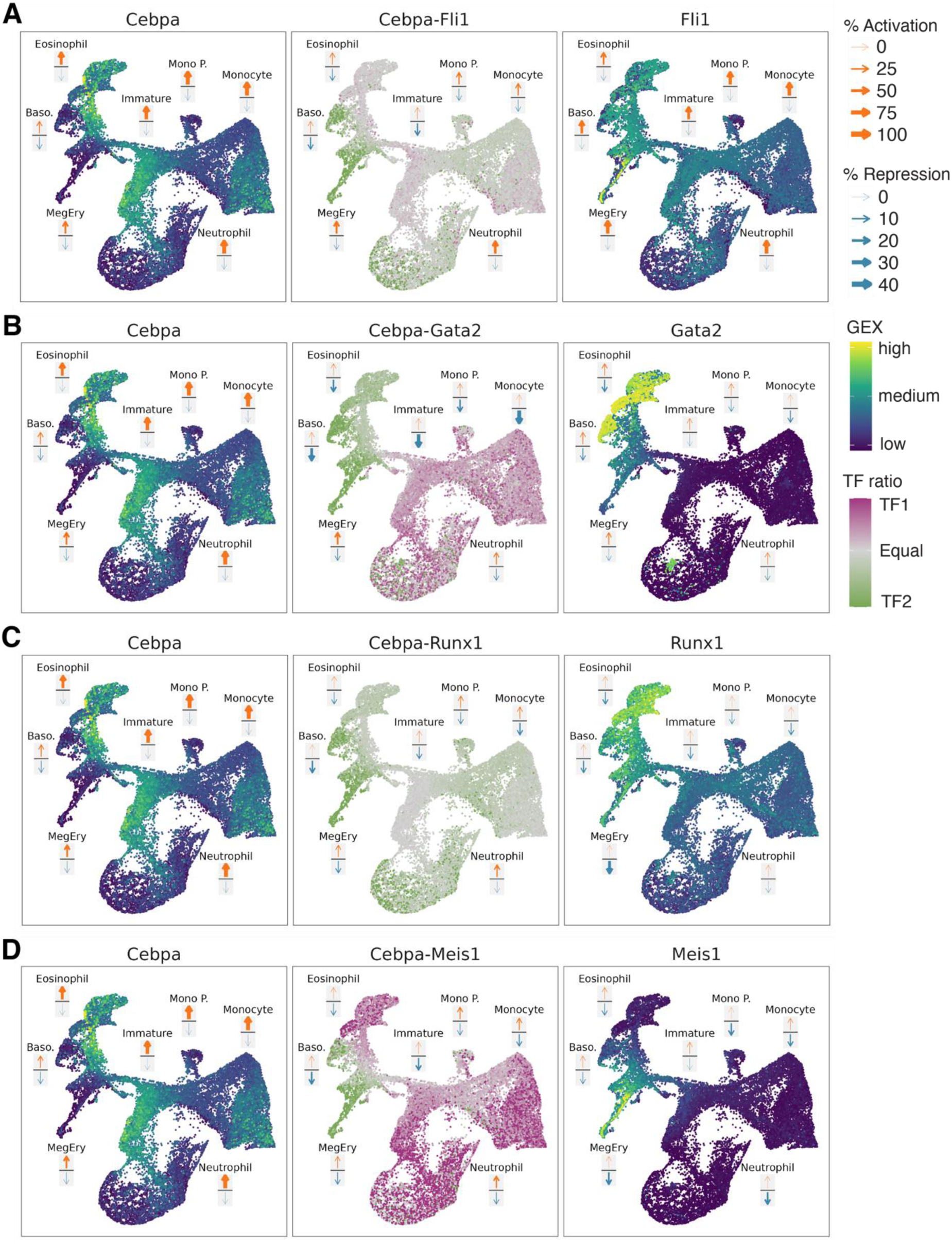
Additional analysis of expression-activity relationships. See main Figure 5c for description.

**Figure S6.**
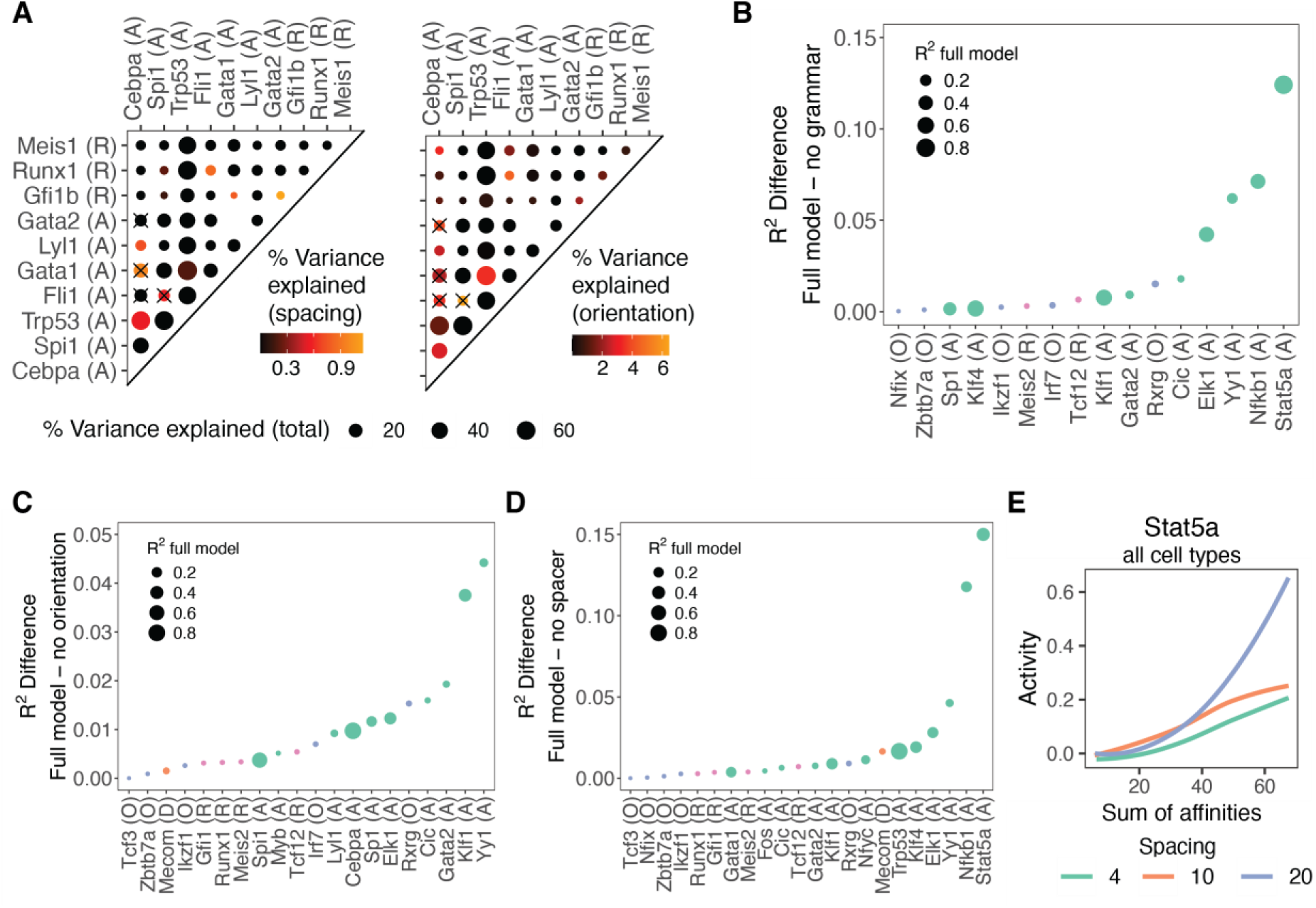
Additional analysis of grammar dependency. See also main Figure 6. **A.** Like main Figure 6a, but contrasting a full model trained on all design features to a model trained on all design features except spacing (left) or orientation (right). **B-D.** Scatter plots depicting, for factors from Library A, the difference in % of variance explained by a random forest trained on all design features, and a model trained on all design features except (b) spacing and orientation, (c) orientation and (d) spacing. Only factors with a non-zero fraction of variance explained by spacing/orientation are shown. Dot size corresponds to performance of the full model. **E.** Line plot displaying smoothened means of activity, conditioned by sum of motif affinities and stratified by spacing between motifs, for Stat5a.

**Figure S7.**
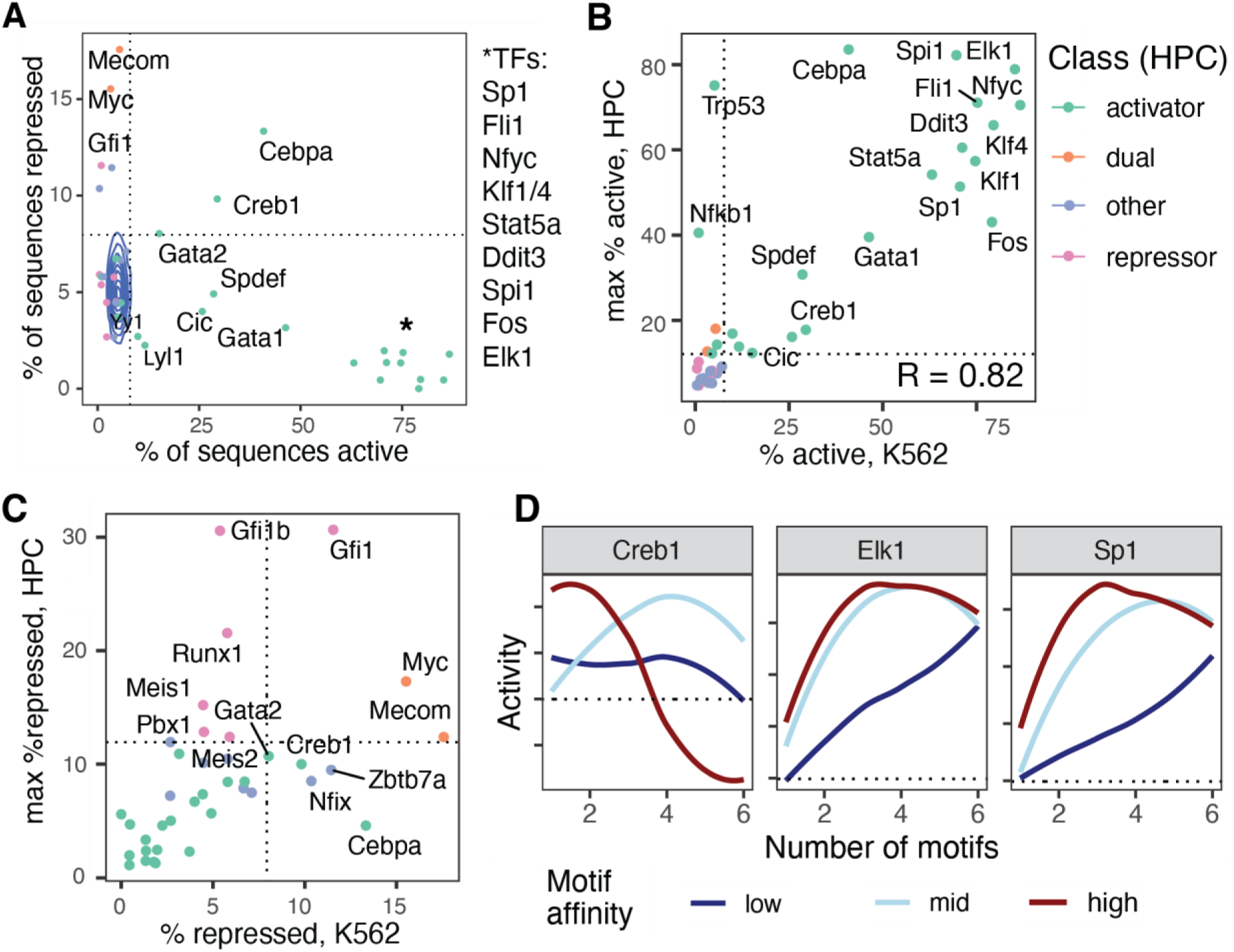
Analysis of single-factor Library A in K562 cells. See Table S1 for source data. **A.** Identification of activators, repressors and dual factors in K562. See main Figure 3a for a legend. Colors correspond to the functional class of the TF identified in HPCs. **B,C.** Scatter plot comparing the ability of the TFs to activate (b) or repress (c) transcription between HPCs and K562 cells. See main Figure 3a for detail. **D.** Dissection of non-monotonous behavior. Line plots displaying smoothened means of activity, conditioned by the number of motifs and stratified by motif affinity, for Creb1, Elk1 and Sp1. See also Figure 4c and S4c.

**Figure S8.**
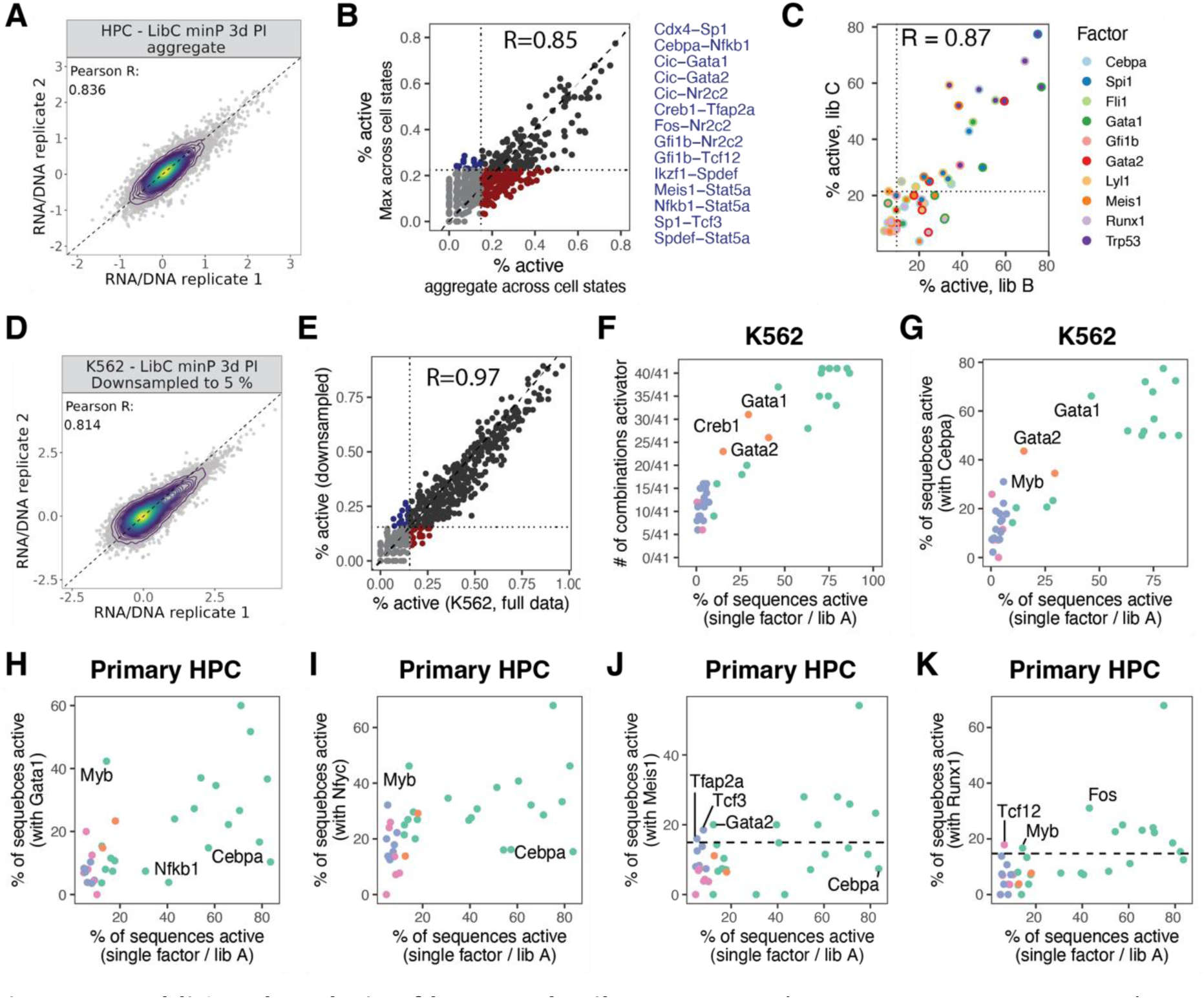
Additional analysis of large-scale Library C. See also main Figure 7. **A.** Correlation between replicates after aggregating primary cell MPRA data across cell states, see also Figure S3. Analysis in main Figure 7 were performed using aggregate data. **B.** Scatter plot comparing, for each pair of TFs, the % of active sequences in aggregated data (x axis), and the maximum % of active sequences across cell states (y axis). Dotted lines indicate the significance threshold for classifying a TF pair as activator, see methods, section *Identification of active and repressed sequences*. Factor pairs identified as activators only in a per-cell-state analysis are listed. **C.** Scatter plot depicting the % of sequences activated by pairs of factors in Library B, and in Library C. Colors of the inner and outer circle denote factor combination. **D.** Correlation between replicates after down-sampling K562 MPRA data to a read depth of 5%. A similar R^2^ compared to the HSPC aggregate (panel A) is obtained. **E.** Scatter plot as in panel B, comparing down-sampled and non down-sampled data from K562 cells. **F.** Scatter plot depicting the relationship between the activity elicited by each single factor in the context of random DNA (x axis) and the number of TF combinations in which the factor acts as an activator (y axis), for K562 cells. **G-K.** Scatter plot depicting the relationship between the activity elicited by a single factor in the context of random DNA (x axis) and the activity elicited by the factor in combination with another factor (y axis), for Cebpa in K562 cells (g), and Gata 1 (h), Nfyc (i), Meis1 (j) and Runx1 (k) in HSPCs. Dashed lines indicate the threshold for classifying a TF pair as activator, see methods, section *Identification of active and repressed sequences*.

**Figure S9.**
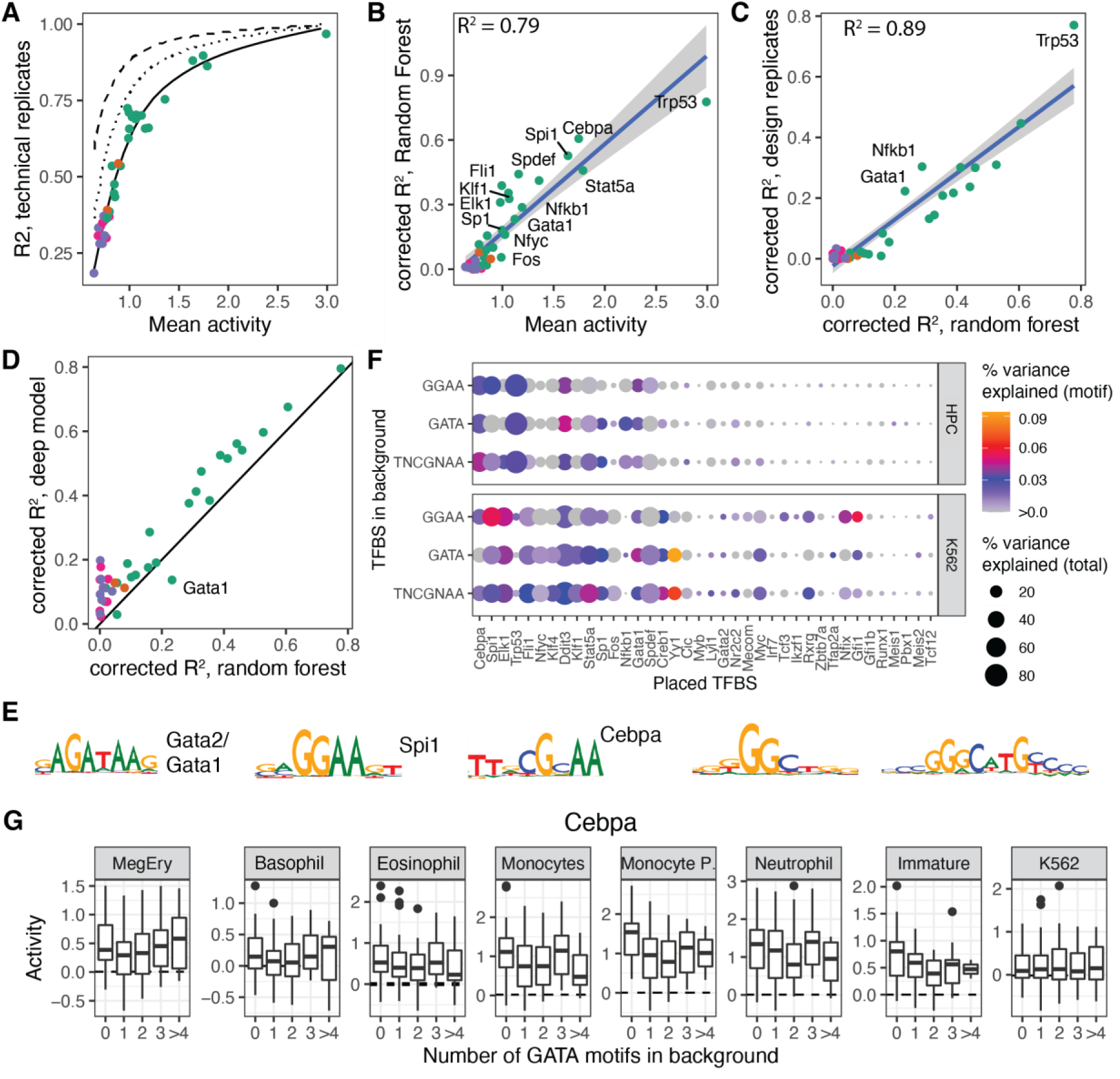
Effects of random background DNA. See also Supplementary Note 2. **A.** Scatter plot depicting, for each transcription factor, the relationship between mean activity and R^2^ between technical replicates. The solid line fits the observed relationship with a monotonously increasing spline function; the dotted line describes the maximum R^2^ that can be achieved between design replicates, given a technical noise level, and the dashed line describes the maximum R^2^ that can be achieved by a machine learning model, given a technical noise level. **B.** Scatter plot depicting the relationship between mean activity and the R^2^ practically achieved by a random forest model during 10-fold cross validation, corrected for the technical noise level. **C.** Scatter plot depicting the relationship between the R^2^ achieved by a random forest model, and the R^2^ between design replicates, in both cases corrected for the technical noise level. **D.** Scatter plot depicting the relationship between the R^2^ achieved by a deep learning model and the R^2^ achieved by a random forest model. **E.** Top five motifs in background DNA sequence with an important impact on deep model predictions, identified by TF-MoDISCO^89^. **F.** Dot plot illustrating the difference in R^2^ between a random forest model trained on design feature, and a random forest model trained on design features plus the number of motif x identified in the background sequence. **G.** Box plot illustrating the effect of GATA motifs in the random DNA background on the activity of sequences containing Cebpa binding sites.

## Supplementary Tables

**Table S1.** Design of libraries.

**Table S2.** Source data for Figure 3a,c and S7a-c.

**Table S3.** Source data for Figure 7.

**Table S4.** Antibodies used for CITE-seq and cell sorting.

**Table S5.** Oligonucleotide sequences

**Table S6.** Data processing parameters

## Supplementary Notes

### Supplementary Note 1: An ex vivo assay of hematopoietic differentiation for massively parallel screens of synthetic cis regulatory elements

We developed an assay to measure the enhancer activity of synthetic DNA throughout hematopoietic differentiation at scale. To that end, we adapted culture conditions^27,29^ that permit the parallel differentiation of primary mouse HSCs into red blood cells, megakaryocytes, neutrophils, eosinophils, basophils, monocytes and dendritic cells. We used single-cell RNA-seq to confirm that the cells produced by this system, as well as their progenitor stages, recapitulate gene expression patterns observed *in vivo* (Figure 1a,b, S2a-d). Specifically, we transferred labels from a single-cell RNA-seq reference of mouse hematopoiesis^75^ (Figure S2a) and additionally annotated cell states based on marker gene expression (Figure S2b,d). We also confirmed that our implementation of these cultures closely reflected the cell state obtained in previous work^27^, although we do obtain a higher fraction of monocytes (Figure S2c).

To stratify these differentiation cultures into discrete progenitor states, we used CITE-seq to design an optimal FACS sorting scheme in a data-driven manner^73^. Specifically, we identified surface antigens specific to the different lineages and used machine learning to identify an optimal gating scheme (Figure S2e-g). Thereby, we stratified the cultures into seven defined cell states after four days of culture, that is, at an intermediate timepoint of differentiation. Residual immature progenitors, megakaryocyte-erythroid progenitors, eosinophil, basophil, neutrophil precursors, as well as early and late monocyte precursors were effectively separated. The purity of the resulting FACS-sorted populations was confirmed by smart-seq2^76^(Figure S2h). While more mature cell states were isolated with purities of >90%, the gradual nature of hematopoiesis makes it impossible to draw hard borders between immature progenitor cell states; still, cells from the same sorting gate clustered together (Figure S2h). We further assessed the reproducibility of our culture conditions by repeatedly isolating RNA from the sorted populations, and performing RNA-seq to demonstrate high reproducibility of our culture conditions (Figure S2i). Overall, these data demonstrate that our primary cell model is a robust and realistic model of hematopoiesis suited for large-scale *ex vivo* MPRA screens.

We then optimized lentiviral MPRA^30,31^ for this setting. In lentiviral MPRA, synthetic DNA is cloned upstream of a minimal promoter and a reporter, and its activity is determined after integration into the genome, i.e. the synthetic DNA is covered in histones. To void effects of chromatin context, the lentiviral vector used is flanked by so-called anti-repressor elements (Figure 1b). Furthermore, the activity measurements of each GRE are averaged over 100s of different integration sites. Activity of a GRE is determined by counting an expressed, GRE-specific barcode in the 5’region of a GFP reporter gene associated at the level of the RNA, and again at the level of DNA, for normalization (see methods).

We cloned the DNA libraries such that each GRE was associated with 100s-1000s of unique barcodes in the 5’UTR of the (Figure S3a). Cloning was very even, leading to a 5-10 fold difference in the representation of the different GREs within the library, with one exception, where the difference was larger (Figure S3b). We then infected primary cells and K562 cells with these libraries and counted the number of barcodes observed per GRE (Figure S3c,d), demonstrating that each element had been integrated at least 50-100 times in primary cells, and 100-1000s of times in K562 cells. Finally, we systematically optimized the computational processing of the MPRA data, using the performance of sequence-to-activity models as an optimization criterion (see methods). We determined that our assay produces high-quality data, as determined by correlations between replicate infections of 0.95-0.99 in the cell line, and 0.74-0.87 in primary cells (Figure S3e,h,i). Taken together, these analyses demonstrate the high technical quality and reproducibility of our measurements.

We performed several additional screens in K562 cells to demonstrate that the choice of core promoter (Figure S3f) or the choice of the measurement timepoint post lentiviral integration (Figure S3g) did not exert a major effect on our results.

### Supplementary Note 2: The effect of random DNA

To evaluate the dependence of sequence activity on the random DNA background, we used Library A, where binding sites for a single factor at a time were placed in a random DNA background. For the sequences of each factor, we compared the fraction of variance in activity explained by design (i.e. number of binding sites, affinity, spacing between sites and site orientation), and the fraction of variance explained by background DNA sequence. We used two approaches, one based on a machine learning model that considers sequence design but not background (“random forest”), and one based on experimental measurements of sequences with similar design, but different background (“design replicates”).

To provide a clean estimate for fraction of variance explained by design and background, it was important to account for the technical noise of the measurements. For factors that drove high levels of expression, the correlation between replicates (technical R^2^) was higher, compared to factors that drove low levels of expression (Figure S9a, solid line). The maximum fraction of variance explained by a machine learning model relates to this technical R^2^, but is higher, since the machine learning model provides an estimate of the expected activity, free of technical noise (Figure S9a, dashed line). The maximum fraction of variance explained by a design replicates also but is higher than the technical R^2^, since here, the correlation between means of replicates is computed (Figure S9a, dotted line).

The analysis of the random forest model demonstrated that Trp53, Spi1 and Cebpa, design explained more than 50% of variance, whereas for the other factors, background dominated (Figure S9b). In general, stronger transcriptional activators were able to dominate over background, even after accounting for the relationship between activity and measurement noise. However, there were factor-specific differences: For example, the activity of constructs containing binding sites for Fos, Nfyc or Sp1 was more strongly impacted by the background sequence, compared to other activators of a similar strength (Fli1, Elk1 or Klf1).

To obtain an experimental validation of these machine-learning based estimates, we made use of “design replicates”, i.e. sequences with a similar sequence design and different background sequences. In the context of Library A, binding site orientation usually had a weak effect on activity (see Figure S6c). We therefore treated sequences with the same number of sites, affinity and spacing between sites as ‘design replicates’ and computed the fraction of variance in one set of design replicates explained by the other design replicate. The estimates of background dependency obtained by this strategy were tightly correlated with the estimates obtained from the machine-learning based strategy (Figure S9c, R^2^ = 0.89).

Finally, we used deep learning models (based on ref. ^90^, implementation available at https://github.com/veltenlab/MPRA_prediction) to identify motifs arising in background sequences that modulate activity. Deep models systematically outperformed random forest model, in particular for factors of low activity where background dominated (Figure S9d). We then used TF-MoDISCO to identify motifs arising in random DNA background that had a strong impact on deep model predictions. The strongest hits were a GATA motif, a GGAA motif resembling the Spi1 and Fli binding sites, and a TTNCGNAA motif resembling the Cebpa binding site (Figure S9e, see also Figure S1c). The presence of these motifs individually explained up to 4% in the variance of activities (Figure S9f). For example, GATA binding sites in the background of sequences containing Cebpa sites weakened the activity of these sequences in primary hematopoietic cells, but not in K562 cells, in line with the results from the combinatorial Library B/C.

Together, these analyses demonstrate that DNA context extensively modulates the activity of hematopoietic TFs; studies that embed transcription factor motifs in a fixed DNA context therefore need to be interpreted with caution.

## References

1. Paul, F. et al. Transcriptional heterogeneity and lineage commitment in myeloid progenitors. Cell 163, 1663–1677 (2015).

2. Velten, L. et al. Human haematopoietic stem cell lineage commitment is a continuous process. Nat. Cell Biol. 19, 271–281 (2017).

3. Tusi, B. K. et al. Population snapshots predict early haematopoietic and erythroid hierarchies. Nature 555, 54–60 (2018).

4. Wheat, J. C. et al. Single-molecule imaging of transcription dynamics in somatic stem cells. Nature (2020) doi:10.1038/s41586-020-2432-4.

5. Loh, K. M. & Lim, B. A precarious balance: Pluripotency factors as lineage specifiers. Cell Stem Cell 8, 363–369 (2011).

6. Pijuan-Sala, B. et al. A single-cell molecular map of mouse gastrulation and early organogenesis. Nature 566, 490–495 (2019).

7. Cockburn, K. et al. Gradual differentiation uncoupled from cell cycle exit generates heterogeneity in the epidermal stem cell layer. Nat. Cell Biol. 24, 1692–1700 (2022).

8. Dulken, B. W., Leeman, D. S., Boutet, S. C., Hebestreit, K. & Brunet, A. Single-Cell Transcriptomic Analysis Defines Heterogeneity and Transcriptional Dynamics in the Adult Neural Stem Cell Lineage. Cell Rep. 18, 777–790 (2017).

9. Dahl, R. et al. Regulation of macrophage and neutrophil cell fates by the PU.1:C/EBPα ratio and granulocyte colony-stimulating factor. Nat. Immunol. 4, 1029–1036 (2003).

10. Hosokawa, H. et al. Transcription Factor PU.1 Represses and Activates Gene Expression in Early T Cells by Redirecting Partner Transcription Factor Binding. Immunity 48, 1119–1134.e7 (2018).

11. Beneyto-Calabuig, S. et al. Clonally resolved single-cell multi-omics identifies routes of cellular differentiation in acute myeloid leukemia. Cell Stem Cell (2023) doi:10.1016/j.stem.2023.04.001.

12. Subramanian, S. et al. Genome-wide transcription factor-binding maps reveal cell-specific changes in the regulatory architecture of human HSPCs. Blood 142, 1448–1462 (2023).

13. Corces, M. R. et al. Lineage-specific and single-cell chromatin accessibility charts human hematopoiesis and leukemia evolution. Nat. Genet. 48, 1193–1203 (2016).

14. Wilson, N. K. et al. Combinatorial Transcriptional Control In Blood Stem/Progenitor Cells: Genome-wide Analysis of Ten Major Transcriptional Regulators. Cell Stem Cell 7, 532–544 (2010).

15. Domingo, J., et al. Non-Linear Transcriptional Responses to Gradual Modulation of Transcription Factor Dosage. http://biorxiv.org/lookup/doi/10.1101/2024.03.01.582837 (2024) doi:10.1101/2024.03.01.582837.

16. Rubin, A. J. et al. Coupled Single-Cell CRISPR Screening and Epigenomic Profiling Reveals Causal Gene Regulatory Networks. Cell 176, 361–376.e17 (2019).

17. White, M. A. et al. A Simple Grammar Defines Activating and Repressing cis-Regulatory Elements in Photoreceptors. Cell Rep. 17, 1247–1254 (2016).

18. Lambert, S. A. et al. The Human Transcription Factors. Cell 172, 650–665 (2018).

19. De Boer, C. G. & Taipale, J. Hold out the genome: a roadmap to solving the cis-regulatory code. Nature (2023) doi:10.1038/s41586-023-06661-w.

20. Fiore, C. & Cohen, B. A. Interactions between pluripotency factors specify cis-regulation in embryonic stem cells. Genome Res. 26, 778–786 (2016).

21. Gertz, J., Siggia, E. D. & Cohen, B. A. Analysis of combinatorial cis-regulation in synthetic and genomic promoters. Nature 457, 215– 218 (2009).

22. Sharon, E. et al. Inferring gene regulatory logic from high-throughput measurements of thousands of systematically designed promoters. Nat. Biotechnol. 30, 521–530 (2012).

23. Georgakopoulos-Soares, I. et al. Transcription factor binding site orientation and order are major drivers of gene regulatory activity. Nat. Commun. 14, 2333 (2023).

24. Levo, M. & Segal, E. In pursuit of design principles of regulatory sequences. Nat. Rev. Genet. 15, 453–468 (2014).

25. de Boer, C. G. et al. Deciphering eukaryotic gene-regulatory logic with 100 million random promoters. Nat. Biotechnol. 38, 56–65 (2020).

26. Sahu, B. et al. Sequence determinants of human gene regulatory elements. Nat. Genet. 54, 283–294 (2022).

27. Weinreb, C., Rodriguez-Fraticelli, A. E., Camargo, F. D. & Klein, A. M. Lineage tracing on transcriptional landscapes links state to fate during differentiation. (2018).

28. Wilkinson, A. C. et al. Long-term ex vivo haematopoietic-stem-cell expansion allows nonconditioned transplantation. Nature (2019) doi:10.1038/s41586-019-1244-x.

29. Jindal, K., et al. Multiomic Single-Cell Lineage Tracing to Dissect Fate-Specific Gene Regulatory Programs. http://biorxiv.org/lookup/doi/10.1101/2022.10.23.512790 (2022) doi:10.1101/2022.10.23.512790.

30. Inoue, F. et al. A systematic comparison reveals substantial differences in chromosomal versus episomal encoding of enhancer activity. Genome Res. 27, 38–52 (2017).

31. Gordon, M. G. et al. lentiMPRA and MPRAflow for high-throughput functional characterization of gene regulatory elements. Nat. Protoc. 15, 2387–2412 (2020).

32. Klein, J. C. et al. A systematic evaluation of the design and context dependencies of massively parallel reporter assays. Nat. Methods (2020) doi:10.1038/s41592-020-0965-y.

33. Agarwal, V., et al. Massively Parallel Characterization of Transcriptional Regulatory Elements in Three Diverse Human Cell Types. http://biorxiv.org/lookup/doi/10.1101/2023.03.05.531189 (2023) doi:10.1101/2023.03.05.531189.

34. Kheradpour, P. et al. Systematic dissection of regulatory motifs in 2000 predicted human enhancers using a massively parallel reporter assay. Genome Res. 23, 800–811 (2013).

35. Zhang, P. et al. Negative cross-talk between hematopoietic regulators: GATA proteins repress PU.1. Proc. Natl. Acad. Sci. U. S. A. 96, 8705–10 (1999).

36. Spooner, C. J., Cheng, J. X., Pujadas, E., Laslo, P. & Singh, H. A recurrent network involving the transcription factors PU.1 and Gfi1 orchestrates innate and adaptive immune cell fates. Immunity 31, 576–86 (2009).

37. Laslo, P. et al. Multilineage Transcriptional Priming and Determination of Alternate Hematopoietic Cell Fates. Cell 126, 755–766 (2006).

38. Pongubala, J. M. R. et al. Transcription factor EBF restricts alternative lineage options and promotes B cell fate commitment independently of Pax5. Nat. Immunol. 9, 203–15 (2008).

39. Friedman, A. D. Transcriptional control of granulocyte and monocyte development. Oncogene 26, 6816–6828 (2007).

40. Ceredig, R., Rolink, A. G. & Brown, G. Models of haematopoiesis: seeing the wood for the trees. Nat. Rev. Immunol. 9, 293–300 (2009).

41. Mercer, E. M., Lin, Y. C. & Murre, C. Factors and networks that underpin early hematopoiesis. Semin. Immunol. 23, 317–25 (2011).

42. Novershtern, N. et al. Densely interconnected transcriptional circuits control cell states in human hematopoiesis. Cell 144, 296–309 (2011).

43. Granja, J. M. et al. Single-cell multiomic analysis identifies regulatory programs in mixed-phenotype acute leukemia. Nat. Biotechnol. 37, 1458–1465 (2019).

44. Kuvardina, O. N. et al. RUNX1 represses the erythroid gene expression program during megakaryocytic differentiation. Blood 125, 3570–3579 (2015).

45. Martinez-Corral, R. et al. Emergence of activation or repression in transcriptional control under a fixed molecular context. Preprint at 10.1101/2024.05.29.596388 (2024).

46. Elagib, K. E. et al. RUNX1 and GATA-1 coexpression and cooperation in megakaryocytic differentiation. Blood 101, 4333–41 (2003).

47. Piras, F. et al. Lentiviral vectors escape innate sensing but trigger p53 in human hematopoietic stem and progenitor cells. EMBO Mol. Med. 9, 1198–1211 (2017).

48. Law, J. C., Ritke, M. K., Yalowich, J. C., Leder, G. H. & Ferrell, R. E. Mutational inactivation of the p53 gene in the human erythroid leukemic K562 cell line. Leuk. Res. 17, 1045–1050 (1993).

49. Lutz, M. et al. Transcriptional repression by the insulator protein CTCF involves histone deacetylases. Nucleic Acids Res. 28, 1707– 1713 (2000).

50. Schmidt, S. F., Larsen, B. D., Loft, A. & Mandrup, S. Cofactor squelching: Artifact or fact? BioEssays 38, 618–626 (2016).

51. Gill, G. & Ptashne, M. Negative effect of the transcriptional activator GAL4. Nature 334, 721–724 (1988).

52. Bell, C. C. et al. Comparative cofactor screens show the influence of transactivation domains and core promoters on the mechanisms of transcription. Nat. Genet. (2024) doi:10.1038/s41588-024-01749-z.

53. Maicas, M. et al. The MDS and EVI1 complex locus (MECOM) isoforms regulate their own transcription and have different roles in the transformation of hematopoietic stem and progenitor cells. Biochim. Biophys. Acta BBA - Gene Regul. Mech. 1860, 721–729 (2017).

54. Pan, D. & Courey, A. J. The same dorsal binding site mediates both activation and repression in a context-dependent manner. EMBO J. 11, 1837–1842 (1992).

55. Mukund, A. X. et al. High-throughput functional characterization of combinations of transcriptional activators and repressors. Cell Syst. 14, 746–763.e5 (2023).

56. Papaemmanuil, E. et al. Genomic Classification and Prognosis in Acute Myeloid Leukemia. N. Engl. J. Med. 374, 2209–2221 (2016).

57. Togasaki, E. et al. Frequent somatic mutations in epigenetic regulators in newly diagnosed chronic myeloid leukemia. Blood Cancer J. 7, e559–e559 (2017).

58. Assi, S. A. et al. Subtype-specific regulatory network rewiring in acute myeloid leukemia. Nat. Genet. 51, 151–162 (2019).

59. Gillespie, M. A. et al. Absolute Quantification of Transcription Factors Reveals Principles of Gene Regulation in Erythropoiesis. Mol. Cell 78, 960–974.e11 (2020).

60. De Almeida, B. P. et al. Targeted design of synthetic enhancers for selected tissues in the Drosophila embryo. Nature (2023) doi:10.1038/s41586-023-06905-9.

61. Taskiran, I. I. et al. Cell-type-directed design of synthetic enhancers. Nature 626, 212–220 (2024).

62. Satpathy, A. T. et al. Massively parallel single-cell chromatin landscapes of human immune cell development and intratumoral T cell exhaustion. Nat. Biotechnol. 37, 925–936 (2019).

63. Kulakovskiy, I. V. et al. HOCOMOCO: Towards a complete collection of transcription factor binding models for human and mouse via large-scale ChIP-Seq analysis. Nucleic Acids Res. 46, D252–D259 (2018).

64. Fornes, O. et al. JASPAR 2020: update of the open-access database of transcription factor binding profiles. Nucleic Acids Res. 48, D87– D92 (2020).

65. Weirauch, M. T. et al. Determination and inference of eukaryotic transcription factor sequence specificity. Cell 158, 1431–1443 (2014).

66. Tan, G. & Lenhard, B. TFBSTools: an R/bioconductor package for transcription factor binding site analysis. Bioinformatics 32, 1555– 1556 (2016).

67. Jean-Marie, B. universalmotif. [object Object] 10.18129/B9.BIOC.UNIVERSALMOTIF (2018).

68. Wagih, O. ggseqlogo: a versatile R package for drawing sequence logos. Bioinformatics 33, 3645–3647 (2017).

69. Gu, Z., Eils, R. & Schlesner, M. Complex heatmaps reveal patterns and correlations in multidimensional genomic data. Bioinformatics 32, 2847–2849 (2016).

70. Grant, C. E., Bailey, T. L. & Noble, W. S. FIMO: scanning for occurrences of a given motif. Bioinformatics 27, 1017–1018 (2011).

71. Tornøe, J., Kusk, P., Johansen, T. E. & Jensen, P. R. Generation of a synthetic mammalian promoter library by modification of sequences spacing transcription factor binding sites. Gene 297, 21–32 (2002).

72. Boroujeni, M. E. & Gardaneh, M. The Superiority of Sucrose Cushion Centrifugation to Ultrafiltration and PEGylation in Generating High-Titer Lentivirus Particles and Transducing Stem Cells with Enhanced Efficiency. Mol. Biotechnol. 60, 185–193 (2018).

73. Triana, S. et al. Single-cell proteo-genomic reference maps of the hematopoietic system enable the purification and massive profiling of precisely defined cell states. Nat. Immunol. (2021) doi:10.1038/s41590-021-01059-0.

74. Kiselev, V. Y., Yiu, A. & Hemberg, M. scmap: projection of single-cell RNA-seq data across data sets. Nat. Methods 15, 359–362 (2018).

75. Baccin, C. et al. Combined single-cell and spatial transcriptomics reveal the molecular, cellular and spatial bone marrow niche organization. Nat. Cell Biol. 22, 38–48 (2020).

76. Picelli, S. et al. Smart-seq2 for sensitive full-length transcriptome profiling in single cells. Nat. Methods 10, 1096–1098 (2013).

77. Velten, L. et al. Identification of leukemic and pre-leukemic stem cells by clonal tracking from single-cell transcriptomics. Nat. Commun. 12, 1366 (2021).

78. Hennig, B. P. et al. Large-Scale Low-Cost NGS Library Preparation Using a Robust Tn5 Purification and Tagmentation Protocol. G3 Bethesda Md 8, 79–89 (2018).

79. Bray, N. L., Pimentel, H., Melsted, P. & Pachter, L. Near-optimal probabilistic RNA-seq quantification. Nat. Biotechnol. 34, 525–527 (2016).

80. Li, H. Aligning sequence reads, clone sequences and assembly contigs with BWA-MEM. (2013).

81. Arnold, C. D. et al. Genome-wide quantitative enhancer activity maps identified by STARR-seq. Science 339, 1–4 (2013).

82. Barr, K. A. & Reinitz, J. A sequence level model of an intact locus predicts the location and function of nonadditive enhancers. PLOS ONE 12, e0180861 (2017).

83. Estrada, J., Wong, F., DePace, A. & Gunawardena, J. Information Integration and Energy Expenditure in Gene Regulation. Cell 166, 234–244 (2016).

84. Martinez-Corral, R. et al. Transcriptional kinetic synergy: A complex landscape revealed by integrating modeling and synthetic biology. Cell Syst. 14, 324–339.e7 (2023).

85. Ma, X., Ezer, D., Navarro, C. & Adryan, B. Reliable scaling of position weight matrices for binding strength comparisons between transcription factors. BMC Bioinformatics 16, 265 (2015).

86. Schep, A. N., Wu, B., Buenrostro, J. D. & Greenleaf, W. J. chromVAR: inferring transcription-factor-associated accessibility from single-cell epigenomic data. Nat. Methods 14, 975–978 (2017).

87. Pliner, H. A. et al. Cicero Predicts cis-Regulatory DNA Interactions from Single-Cell Chromatin Accessibility Data. Mol. Cell 71, 858–871.e8 (2018).

88. Li, C. et al. A small regulatory element from chromosome 19 enhances liver-specific gene expression. Gene Ther. 16, 43–51 (2009).

89. Shrikumar, A. et al. Technical Note on Transcription Factor Motif Discovery from Importance Scores (TF-MoDISco) version 0.5.6.5. Preprint at http://arxiv.org/abs/1811.00416 (2020).

90. Janssens, J. et al. Decoding gene regulation in the fly brain. Nature 601, 1–7 (2022).

